# Transposable element and host silencing activity in gigantic genomes

**DOI:** 10.1101/2022.12.20.521252

**Authors:** Jie Wang, Liang Yuan, Jiaxing Tang, Jiongyu Liu, Cheng Sun, Michael W. Itgen, Guiying Chen, Stanley K. Sessions, Guangpu Zhang, Rachel Lockridge Mueller

## Abstract

Transposable elements (TEs) and the silencing machinery of their hosts are engaged in a germline arms-race dynamic that shapes TE accumulation and, therefore, genome size. In animal species with extremely large genomes (>10 Gb), TE accumulation has been pushed to the extreme, prompting the question of whether TE silencing also deviates from typical conditions. To address this question, we characterize TE silencing via two pathways — the piRNA pathway and KRAB-ZFP transcriptional repression — in the male and female gonads of *Ranodon sibiricus*, a salamander species with a ∼21 Gb genome. We quantify 1) genomic TE diversity, 2) TE expression, and 3) small RNA expression and find a significant relationship between the expression of piRNAs and TEs they target for silencing in both sexes. We also quantified TE silencing pathway gene expression in *R. sibiricus* and 14 other vertebrates with genome sizes ranging from 1 – 130 Gb and find no association between pathway expression and genome size. Taken together, our results reveal that the gigantic *R. sibiricus* genome includes at least 19 putatively active TE superfamilies, all of which are targeted by the piRNA pathway in proportion to their expression levels, suggesting comprehensive piRNA-mediated silencing. Males have higher TE expression than females, suggesting that they may contribute more to the species’ high genomic TE load. We posit that apparently conflicting interpretations of TE silencing and genomic gigantism in the literature, as well as the absence of a correlation between TE silencing pathway gene expression and genome size, can be reconciled by considering whether the TE community or the host is currently “on the attack” in the arms race dynamic.

## 1 Introduction

Transposable elements (TEs) are DNA sequences that can mobilize throughout the genomes of their hosts, typically replicating as part of the transposition life cycle (Doolittle and Sapienza, 1980;Orgel and Crick, 1980;Wicker et al., 2007). TEs are an ancient and diverse class of sequences, encompassing a range of replication mechanisms that rely on both TE- and host-encoded enzymatic machinery (Bourque et al., 2018). Eukaryotic genomes contain a substantial yet variable number of TEs; they make up well over half of the human genome, up to 85% of the maize genome (Haberer et al., 2005), yet only ∼0.1% of the yeast *Pseudozyma antarctica* genome (de Koning et al., 2011;Castanera et al., 2017;Jiao et al., 2017). TE abundance is one of the major determinants of overall genome size, which ranges from ∼0.002 Gb to ∼150 Gb across eukaryotes and from ∼0.4 Gb to ∼130 Gb across vertebrates (Rodriguez and Arkhipova, 2018;Gregory, 2022). The mechanistic and evolutionary forces shaping TE abundance and, thus, genome size remain incompletely understood.

Individual TE insertions have a range of effects on host fitness; the majority are effectively neutral or slightly deleterious, while smaller proportions are either harmful (or lethal) on the one hand or adaptive on the other (Arkhipova, 2018;Almeida et al., 2022). For example, at least 120 human diseases have been attributed to the effects of de novo TE insertions, but so have classic adaptive traits including industrial melanism and the mammalian placenta (Hancks and Kazazian, 2016;Hof et al., 2016;Senft and Macfarlan, 2021). The likelihood of a novel TE insertion having an extreme effect on the host phenotype depends on properties of the host genome including gene density, which affects the probability of a new insertion disrupting a functional protein-coding or regulatory sequence (Medstrand et al., 2002).

In response to TEs’ mutagenic properties, eukaryotes have evolved multiple mechanisms to silence their activity, particularly in the germline and early embryo where TE effects on host fitness are the most pronounced (Almeida et al., 2022). Some mechanisms act by transcriptionally silencing TE loci through targeted deposition of chromatin modifications (e.g. methylation of cytosines on DNA, or H3K9 methylation of histone proteins) (Deniz et al., 2019). Other mechanisms act post-transcriptionally, targeting TE transcripts for destruction in the cytoplasm before they can complete the replicative life cycle and generate a novel genomic TE insertion (Czech and Hannon, 2016).

In multicellular animals, TE silencing in the germline and during early embryogenesis is carried out by the piRNA pathway, a small RNA pathway that relies on RNA-induced silencing complexes (RISC) composed of PIWI proteins and associated guide piRNAs that identify TE transcripts by base complementarity (Aravin et al., 2006;Ozata et al., 2019;Iwakawa and Tomari, 2022). In the nucleus, piRNA-PIWI complexes identify chromatin-associated nascent TE transcripts, inducing transcriptional silencing of the genomic TE locus through recruitment of DNA methyltransferases and histone methyltransferases to establish a repressive chromatin structure (Aravin et al., 2008;Czech et al., 2018). In the cytoplasm, piRNA-PIWI complexes identify mature TE transcripts and cleave them between nucleotide positions 10 and 11 of the guide piRNA (Reuter et al., 2011;Iwasaki et al., 2015). The cleaved fragments of TE mRNA induce the production of more TE-targeting piRNAs through a feed-forward loop called the ping-pong cycle, which amplifies the cell’s post-transcriptional TE silencing response (Brennecke et al., 2007;Gunawardane et al., 2007;Castel and Martienssen, 2013). Although present in both males and females, there are sex-specific differences in activity of the piRNA pathway, which may be associated with sex-biased contributions to overall genomic TE load (Saint-Leandre et al., 2020).

In lobe-finned fishes (tetrapods, coelacanth, and lungfishes), TE silencing is also carried out by a large family of transcriptional modulators called the Krüppel-associated box domain-containing zinc-finger proteins (KRAB-ZFPs) (Imbeault et al., 2017). These proteins include an array of zinc fingers, each of which binds short DNA sequences such that, together, they confer specificity to individual TE families (Thomas and Schneider, 2011). These proteins also include the KRAB domain, which recruits KAP1/TRIM28 and, in turn, a silencing complex of proteins that establish a repressive chromatin structure at TE loci (Ecco et al., 2017).

Although these TE silencing pathways are broadly conserved phylogenetically and functionally critical for maintaining genome integrity, they nonetheless evolve (Parhad and Theurkauf, 2019). Our work is motivated by the hypothesis that their evolution contributes to variation in TE content, and therefore overall genome size, across the tree of life (Mueller, 2017). Species that are extreme genome size outliers provide a powerful test of this hypothesis, as they are predicted to harbor strong signatures of divergent TE silencing compared with genomes of more typical size.

Among vertebrates, extreme genome expansion through TE accumulation evolved independently in salamanders and in lungfishes, with large increases in both lineages occurring over 200 million years ago (Liedtke et al., 2018;Meyer et al., 2021). Salamanders are one of the three clades of living amphibians; there are 775 extant species, and haploid genome sizes range from 9 to 120 Gb, reflecting ongoing genome size evolution (AmphibiaWeb, 2022;Gregory, 2022). Lungfishes are the sister taxon to tetrapods; there are 6 extant species, and haploid genome size estimates range from 40 Gb to 130 Gb (Meyer et al., 2021). Amphibians also include some of the smallest vertebrate genomes; the ornate burrowing frog *Platyplectrum ornatum* and the New Mexico spadefoot toad *Spea multiplicata* have genome sizes of 1.06 and 1.09 Gb, respectively (Lamichhaney et al., 2021;Gregory, 2022).

To date, several studies have begun to explore the relationship between TE silencing and genome size among vertebrates. At the smaller extremes, studies of frogs and fish with tiny genomes (≤ 1 Gb) revealed at least one additional duplicate copy of a PIWI gene, suggesting increased activity of the piRNA pathway in silencing TEs in genomes that have undergone size reduction (Malmstrøm et al., 2018;Lamichhaney et al., 2021). At the larger extremes, the data reveal a more complex picture; the Australian and African lungfish genomes (*Neoceratodus forsteri* and *Protopterus annectens*, ≥ 40 Gb) show neither gains nor losses of PIWI or related genes (Biscotti et al., 2017;Meyer et al., 2021). However, the African lungfish genome includes far more KRAB domains than other vertebrate genomes, suggesting a copy-number-based increase in activity. In contrast, the genome of the Mexican axolotl salamander *Ambystoma mexicanum* (∼34 Gb) (Gregory, 2022) contains a comparable number of KRAB domains to mammalian and non-avian reptile genomes, suggesting no similar increase in this TE silencing activity (Wang et al., 2021b).

Expression data reveal a similarly mixed picture: for some piRNA pathway genes, germline expression is higher in salamanders (represented by the fire-bellied newt *Cynops orientalis*, ∼43 Gb) than in the African lungfish, whereas for other genes, the pattern is reversed; comparisons with genomes of more typical size (coelacanth *Latimeria menadoensis* and zebrafish *Danio rerio*) show patterns of both higher and lower germline expression of TE silencing genes in the species with gigantic genomes (Biscotti et al., 2017;Carducci et al., 2021). Small RNA sequence data from the gonads of the northern dusky salamander *Desmognathus fuscus* (∼15 Gb) reveal lower percentages of TE-mapping piRNAs than are found in smaller genomes, suggesting a less comprehensive TE-targeting piRNA pool in the gigantic genome (Madison-Villar et al., 2016). Taken together, these inconsistent patterns reveal that the relationship between TE silencing pathway activity and genome size evolution remains incompletely understood, and that integrating genomic, transcriptomic, and small RNA analysis is critical for a complete picture.

Here we present a detailed analysis of TEs and germline TE silencing activity in the central Asian salamander *Ranodon sibiricus* — a range-restricted species endemic to China and Kazakhstan — adding both phylogenetic (family Hynobiidae) and genome size (∼21 Gb) diversity to the small but growing dataset on TE silencing in gigantic genomes (AmphibiaWeb, 2022;Gregory, 2022). We quantify the expression of TEs in the male and female gonads, and we complement this data with analyses of the genomic TE landscape and TE amplification histories to reveal what TE superfamilies are active in the *R. sibiricus* genome. We quantify small RNAs expressed in male and female gonads and test whether small RNAs targeting TEs for silencing are expressed and amplified in proportion to TE expression. We quantify the relative expression of genes encoding proteins from two TE silencing pathways — piRNA and KRAB-ZFP. Finally, we extend these latter analyses to other vertebrates with a range of genome sizes to test for changes in TE silencing accompanying extreme increases in genome size.

## 2 Results

### 2.1 The genome of *R. sibiricus* contains diverse known, active TE superfamilies

We estimated the haploid genome size of *R. sibiricus* to be 17 Gb; averaging our result with published estimates (22.3 or 24.8 Gb) yields 21.3 Gb (Gregory, 2022). We used the PiRATE pipeline (Berthelier et al., 2018), which was designed to mine and classify repeats from low-coverage genomic shotgun data in taxa that lack genomic resources. The pipeline yielded 109,909 repeat contigs (Table1). RepeatMasker mined the most repeats (75,381 out of 109,909; 68.6%), followed by dnaPipeTE (21.9%), RepeatScout (3.3%), RepeatModeler (2.9%), and TE-HMMER (2.8%). TEdenovo, LTRharvest, HelSearch, SINE-Finder, and MITE-Hunter found few repeats, and MGEScan-non-LTR found none.

**Table 1.**
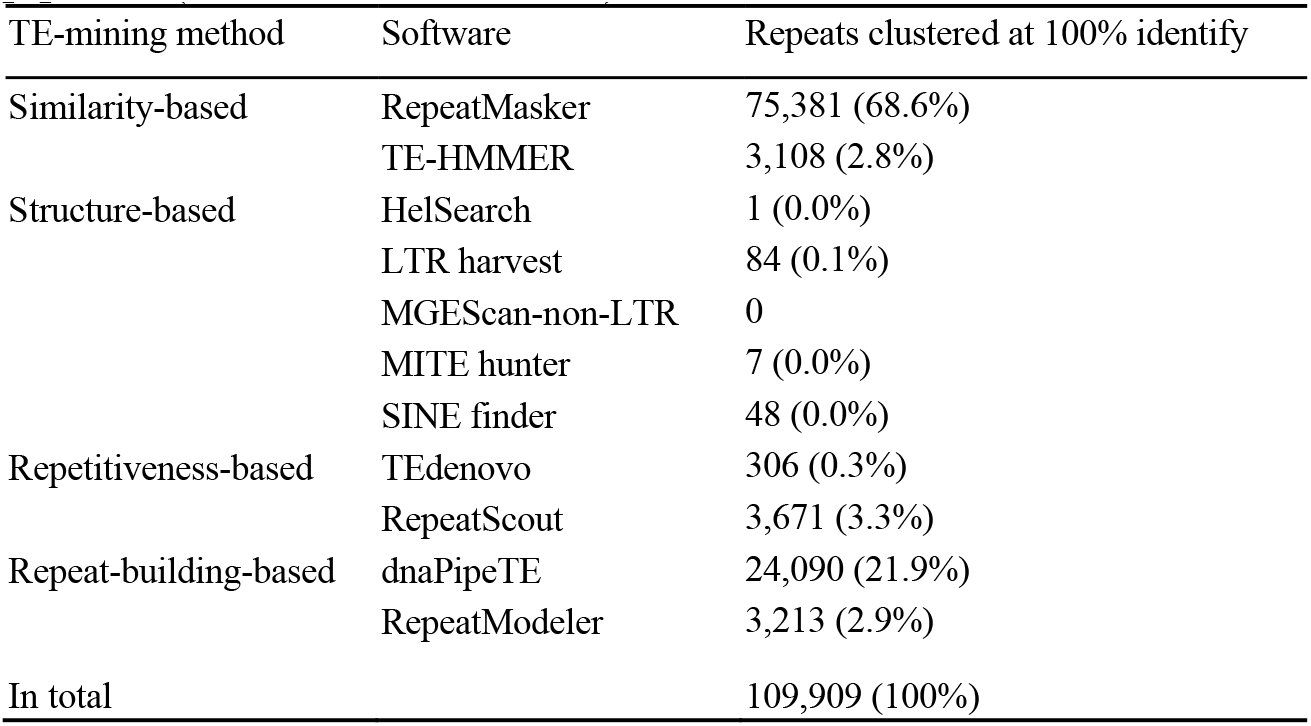
Repeat contigs (≥ 100 bp) identified by different methods/software in the PiRATE pipeline (Berthelier et al., 2018).

Repeat contigs were annotated as TEs to the levels of order and superfamily in Wicker’s hierarchical classification system (Wicker et al., 2007), modified to include several recently discovered TE superfamilies, using PASTEC (Hoede et al., 2014). Of the 109,909 identified repeat contigs, 1,088 were filtered out as potential chimeras, 275 were classified as potential multiple-copy host genes, and 54,221 (49.33%) were classified as known TEs (Table 2), representing 23 superfamilies in eight orders as well as retrotransposon and transposon derivatives.

**Table 2.** Classification of repeat contigs (modified from Wicker 2007) and summary of repeats detected in the genome.

| Order | Superfamily | Percent of Genome<sup>a</sup> | Genomic Contigs (100% Identical) | Genomic Contigs (95% Identical) | Genomic Contigs (80% Identical) | Average Genomic Contig Length (100% identical) (bp) | Longest Genomic Contig (bp) | Transcriptome contigs (80% Identical) | Expression Level in females (TPM) | Expression Level in males (TPM) |
| --- | --- | --- | --- | --- | --- | --- | --- | --- | --- | --- |
| **Class I - Retrotransposons - Autonomous** |  |  |  |  |  |  |  |  |  |  |
| LTR | *Gypsy* | 3.85-6.50 | 5,985 | 5,464 | 3,835 | 548 | 7,587 | 2,057 | 827 | 3,024 |
|  | *ERV* | 0.41-0.43 | 413 | 394 | 264 | 593 | 11,567 | 321 | 392 | 694 |
|  | *Copia* | 0.10-0.25 | 23 | 21 | 19 | 592 | 1,353 | 27 | 31 | 26 |
|  | *Bel-Pao* | 0.05 | 12 | 7 | 2 | 575 | 766 | - | - | - |
|  | *Retrovirus* | - | 3 | 2 | 2 | 1,375 | 1,719 | 10 | 1 | 10 |
|  | *THE1* | - | 4 | 4 | 3 | 491 | 674 | - | - | - |
|  | *Unknown LTR* | - | 4 | 4 | 4 | 469 | 955 | - | - | - |
| DIRS | *DIRS* | 4.44-5.95 | 8,087 | 7,821 | 3,844 | 379 | 4,266 | 4,844 | 5,427 | 11,962 |
| PLE | *Penelope* | 0.09-0.12 | 482 | 462 | 376 | 407 | 3,208 | 460 | 123 | 352 |
| LINE | *Jockey* | 9.69-12.12 | 25,276 | 23,361 | 13,287 | 426 | 3,625 | 11,622 | 6,206 | 15,535 |
|  | *L1* | 5.04-6.62 | 10,189 | 8,997 | 5,941 | 672 | 6,546 | 8,241 | 2,954 | 7,610 |
|  | *RTE* | 0.12-0.24 | 227 | 201 | 137 | 676 | 4,540 | 399 | 130 | 360 |
|  | *I* | 0.09-0.17 | 32 | 25 | 10 | 905 | 3,858 | 48 | 42 | 125 |
|  | *R2* | - |  |  |  |  |  | 1 | - | 1 |
|  | *Unknown LINE* | 0.39-1.89 | 90 | 77 | 72 | 1,393 | 5,881 | 1 | - | - |
| **Class I - Retrotransposons - Non-autonomous** |  |  |  |  |  |  |  |  |  |  |
| SINE | *5S* | 0.23-0.18 | 57 | 41 | 15 | 773 | 1,783 | 4 | 5 | 1 |
|  | *7SL* | - | 43 | 41 | 15 | 243 | 450 | 2 | 1 | - |
|  | *tRNA* | - | 22 | 22 | 22 | 273 | 403 | - | - | - |
|  | *Unknown SINE* | 0.45-3.42 | 365 | 348 | 338 | 377 | 695 | 219 | 2,901 | 2,471 |
| Retrotransposon Derivatives | TRIM | 3.80-9.76 | 748 | 631 | 465 | 588 | 2,619 | 492 | 1,651 | 5,847 |
|  | LARD | 0.15-0.73 | 45 | 37 | 35 | 2,834 | 8,148 | 153 | 515 | 2,643 |
| **Class II - DNA Transposons - Subclass 1** |  |  |  |  |  |  |  |  |  |  |
| TIR | *PIF-Harbinger* | 2.98-4.22 | 1,164 | 1,014 | 434 | 411 | 5,169 | 416 | 575 | 1,359 |
|  | *hAT* | 1.15-1.12 | 177 | 135 | 55 | 869 | 5,706 | 35 | 83 | 26 |
|  | *Tc1-Mariner* | 0.18-0.63 | 73 | 40 | 24 | 1025 | 3,781 | 20 | 58 | 87 |
|  | *PiggyBac* | 0.05-0.08 | 9 | 8 | 6 | 1,087 | 1,956 | 6 | 21 | 15 |
|  | *MuDR* | - | 3 | 3 | 3 | 266 | 417 | 13 | 4 | 10 |
|  | *CACTA* | - | 2 | 2 | 2 | 645 | 846 | 3 | 2 | 1 |
|  | *ISL2EU* | - |  |  |  |  |  | 1 | 18 | 16 |
|  | *Ginger* | - |  |  |  |  |  | 4 | 10 | 6 |
|  | *Academ* | - |  |  |  |  |  | 11 | 5 | 8 |
|  | *P* | - |  |  |  |  |  | 1 | 1 | 1 |
|  | *Unknown TIR* | 0.33-1.46 | 22 | 20 | 19 | 754 | 2,174 | - | - | - |
| Transposon Derivatives | MITE | 0.56-4.07 | 357 | 346 | 321 | 258 | 1,942 | 137 | 465 | 1,188 |
| **Class II - DNA Transposons - Subclass 2** |  |  |  |  |  |  |  |  |  |  |
| Maverick | *Maverick* | 0.05 | 258 | 228 | 65 | 583 | 6,090 | 123 | 44 | 762 |
| Helitron | *Helitron* | 0.18-0.48 | 49 | 38 | 19 | 939 | 6,551 | 13 | 2 | 10 |
| **Total** |  | 34.38-60.54 | 54,221 | 49,794 | 29,634 | 489 | 11,567 | 29,684 | 22,494 | 42,200 |

<sup>a</sup>The first and second numbers were estimated including and excluding unknown repeats, respectively, from the repeat library.

To calculate the proportion of different repeats in the genome, shotgun reads were masked with RepeatMasker using two *R. sibiricus*-derived repeat libraries: excluding or including unknown repeats. This comparison provided a rough approximation of the quantity of unknown repeats that were TE-derived, but divergent, fragmented, or otherwise unidentifiable by our pipeline.

Class I TEs (retrotransposons) make up 28.90%–48.43% (unknown repeats included or excluded in the repeat library, respectively) of the *R. sibiricus* genome; they are over 4 times more abundant than Class II TEs (DNA transposons; 5.48%–12.11%). *LINE/Jockey* is the most abundant superfamily (9.69%–12.12% of the genome), followed by *LINE/L1* (5.04%–6.62%), *LTR/Gypsy* (3.85–6.50%), and TRIM (3.80%–9.76%); all are retrotransposons or retrotransposon derivatives (Table 2). *TIR/PIF-harbinger* (2.98%– 4.22%), *TIR/hAT* (1.12%–1.15%), and MITE (0.56%–4.07%) are the most abundant superfamilies of DNA transposons/transposon derivatives (Table 2).

Diversity of the overall genomic TE community was measured using both Simpson’s and Shannon diversity indices, considering TE superfamilies as “species” and the total number of base pairs for each annotated superfamily as individuals per “species.” The Gini-Simpson Index (1-D) is 0.83, and the Shannon Index H is 1.92, similar to estimates of genomic diversity from other salamander species (Wang et al., 2021a;Haley and Mueller, 2022).

Seventeen superfamilies and three retrotransposon or transposon derivatives (each covering more than 0.05% of the genome) were selected for summaries of overall amplification history, generated by plotting the genetic distances between individual reads (representing TE loci) and the corresponding ancestral TE sequences as a histogram, with bins of 1%. All of the resulting distributions showed characteristics of ongoing or recent activity (*i*.*e*., presence of TE sequences < 1% diverged from the ancestral sequence) (Figure 1).

**Figure 1.**
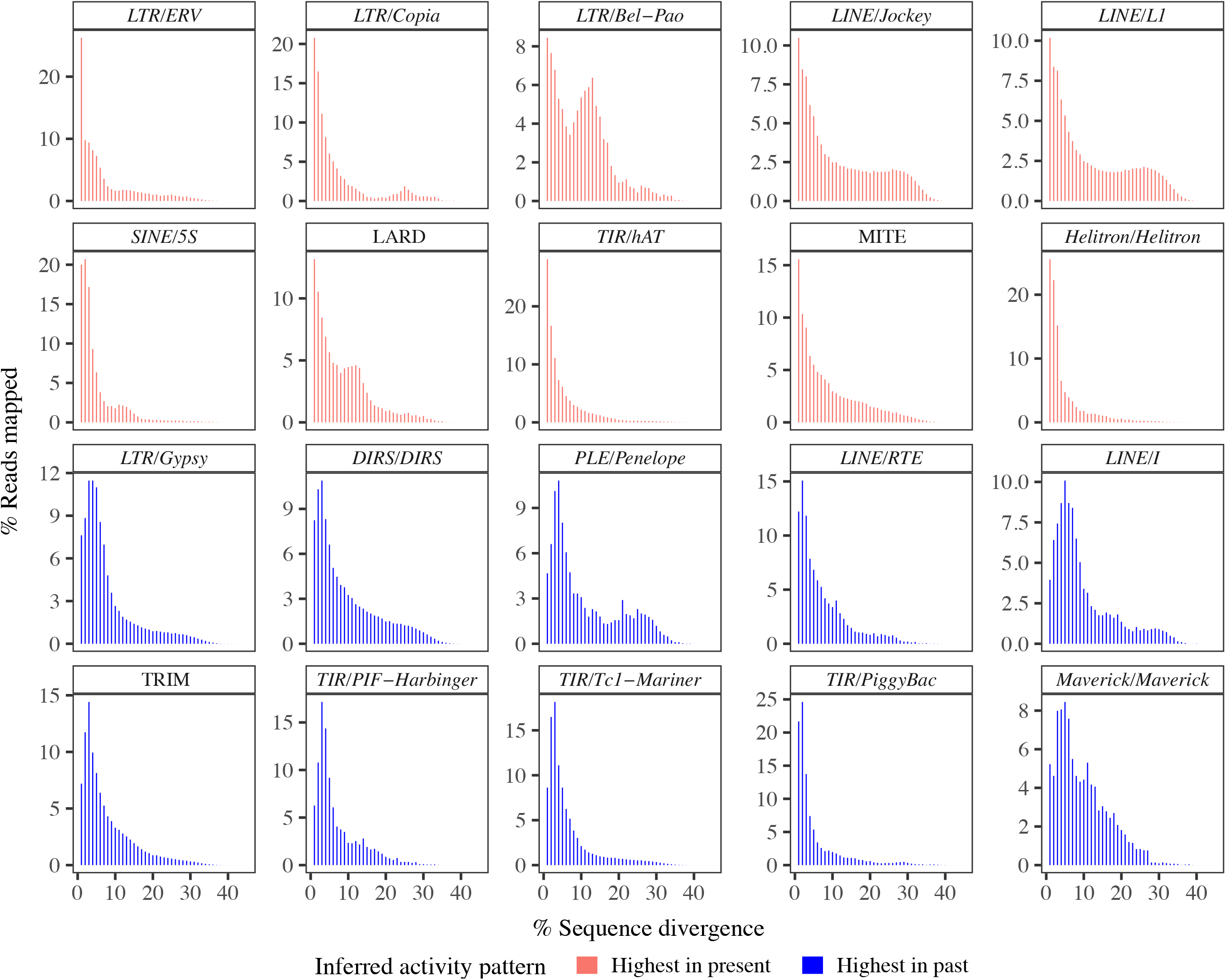
Amplification plots for TE superfamilies and derivatives. All of the amplification plots suggest current activity. Note that the y-axes differ in scale.

Ten of these showed right-skewed, essentially monotonically decreasing distributions with a maximum or near-maximum at < 1% diverged from the ancestral sequence: *LTR/ERV, LTR/Copia, LTR/Bel-Pao, LINE/Jockey, LINE/L1, SINE/5S*, LARD, *TIR/hAT*, MITE, and *Helitron/Helitron*, suggesting TE superfamilies or derivatives that continue to be replicating today at their highest-ever rates of accumulation. In contrast, ten TE superfamilies or derivatives showed right-skewed, uni- or multimodal distributions: *LTR/Gypsy, DIRS/DIRS, PLE/Penelope, LINE/I, LINE/RTE*, TRIM, *TIR/PIF-Harbinger, TIR/Tc1-Mariner, TIR/PiggyBac*, and *Maverick/Maverick*. These 10 distributions suggest TE superfamilies that continue to be active today, but whose accumulation peaked at some point in the past.

### 2.2 Germline relative TE expression is higher in males than females, but is correlated with genomic abundance in both sexes

Our *de novo* gonad transcriptome assembly yielded 510,439 contigs (N50 = 1,250 bp; min and max contig lengths = 201 bp and 28,590 bp; total assembly length = 362,097,394 bp). The BUSCO pipeline revealed the presence of 95.3% of core vertebrate genes and 89.8% of core tetrapod genes. 47,182 contigs were annotated as TEs (representing 28 superfamilies), 64,409 as endogenous genes (representing 28,283 different genes), and 1,257 as having both a TE and an endogenous gene; the majority of contigs (72%) remained unannotated.

Endogenous genes account for the majority of expression in the gonads of both sexes (68% and 51% of summed TPM in females and males, respectively), followed by unannotated contigs (29% and 42%). Relative expression of TEs is an order of magnitude lower than endogenous gene expression in both sexes (2.4% and 5.6%) (Table 3, Supplementary File S3). Nine superfamilies (*LTR/Retrovirus, LINE/R2, SINE/7SL, TIR/MuDR, TIR/CACTA, TIR/ISL2EU, TIR/Ginger, TIR/Academ, TIR/P*) were detected at low expression levels in the transcriptome but were not initially detected in the genomic data (Table 2); mapping the genomic reads to these transcriptome contigs with Bowtie2 identified an average of 4 reads per superfamily, indicating their extremely low frequency in the genome. In contrast, only one superfamily (*LTR/Bel-Pao*) was detected in the genomic data but not in the transcriptome data. Overall, 19 superfamilies were identified both in the genomic contigs and transcriptome contigs.

**Table 3.** Overall summary of transcriptome annotation and expression in each sex.

|  | No. of contigs expressed in females (percentage of total contigs) | Summed TPM in females (percentage of total expression) | No. of contigs expressed in males (percentage of total contigs) | Summed TPM in males (percentage of total expression) |
| --- | --- | --- | --- | --- |
| Endogenous gene | 51,647 (17.4%) | 678,366 (67.8%) | 58,041 (13.3%) | 513,783 (51.4%) |
| Autonomous TE | 26,358 (8.9%) | 17,890 (1.8%) | 41,421 (9.5%) | 43,694 (4.4%) |
| Non-autonomous TE | 1,700 (0.6%) | 5,538 (0.6%) | 2,348 (0.5%) | 12,151 (1.2%) |
| Gene/autonomous TE | 722 (0.2%) | 2,618 (0.3%) | 1,001 (0.2%) | 5,748 (0.6%) |
| Gene/nonautonomous TE | 168 (0.1%) | 2,779 (0.3%) | 189 (0.0%) | 1,073 (0.1%) |
| Unannotated | 215,508 (72.8%) | 292,828 (29.3%) | 334,533 (76.5%) | 423,550 (42.4%) |
| Total | 296,103 (100%) | 1,000,029 (100%) | 437,533 (100%) | 999,999 (100%) |

In ovaries, autonomous TEs account for 8.9% of the total transcriptome contigs and 1.8% of the overall transcripts (summed TPM = 17,890) (Table 3). Non-autonomous TEs account for only 0.6% of the total transcriptome contigs, but still represent 0.6% of the overall transcripts (summed TPM = 5,538). In testes, relative TE expression is more than double that seen in females, with 4.4% and 1.2% of the overall gonad transcriptome accounted for by autonomous and non-autonomous TEs, respectively.

Differential expression analysis identified 780 contigs of 18 TE superfamilies as differently expressed between testes and ovaries. 678 TE transcripts were more highly expressed in testes, while only 102 TE transcripts were more highly expressed in ovaries (Figure 2A). Of the nine superfamilies with more than ten differentially expressed transcripts between males and females, eight of them showed significantly higher relative expression in testes than ovaries, and one showed a non-significant trend towards higher testis expression (Figure 2B). Across the 19 TE superfamilies detected in both the genomic and transcriptomic datasets, genomic abundance is positively correlated with overall relative expression both in ovaries (R = 0.786, P < 0.001) and testes (R=0.837, P < 0.001), with male relative expression higher overall (Figure 2C).

**Figure 2.**
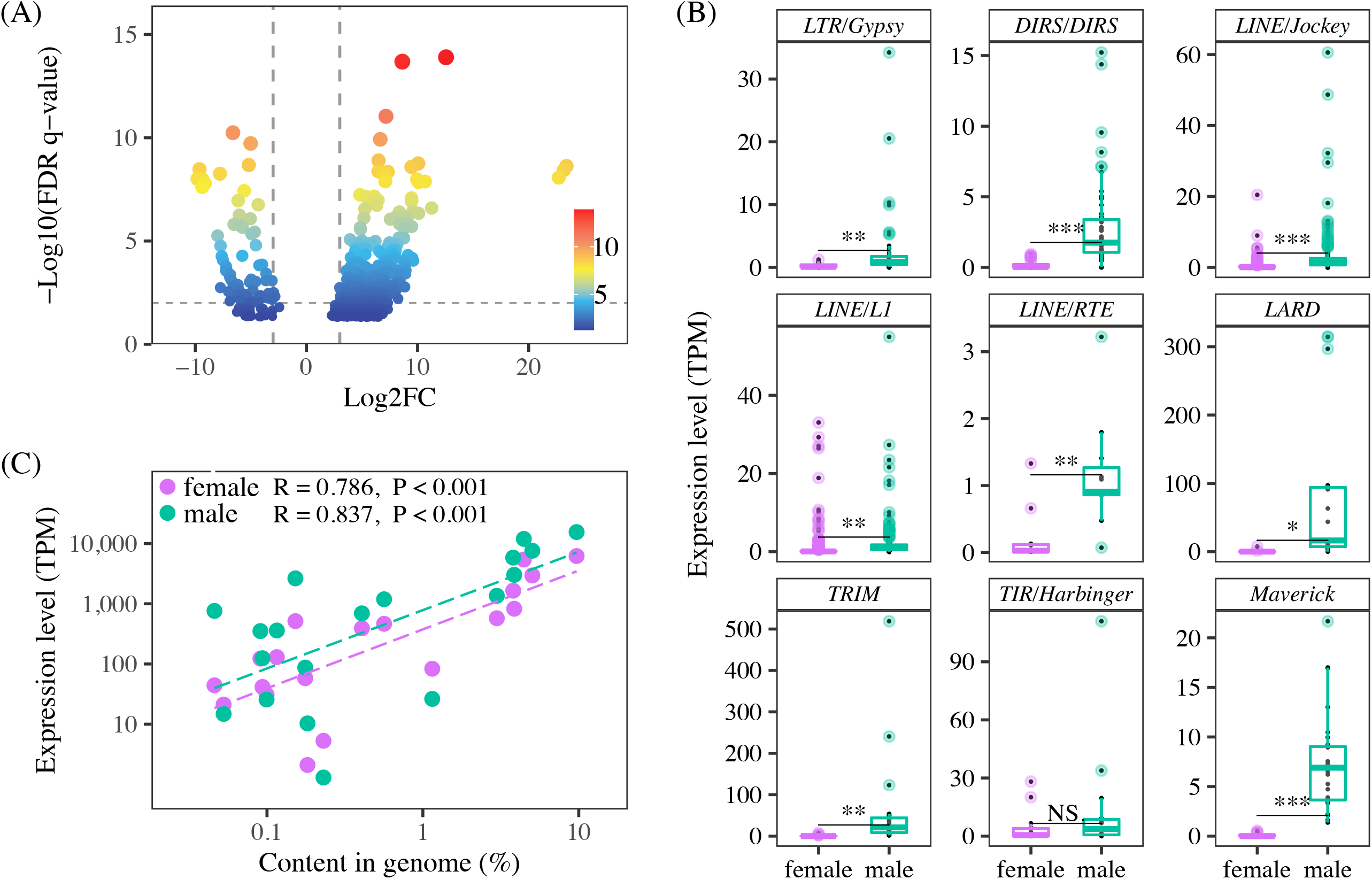
(A) TE transcripts that are differentially expressed between testes and ovaries. Positive fold-change value indicates higher expression in testes. The majority of transcripts show male-biased expression. (B) Expression levels of TE superfamilies represented by >10 TE transcripts in ovaries vs. testes. Expression is higher in males. Each point represents the average expression of a TE transcript across same-sex individuals. * and NS indicate P < 0.05 and not significant, respectively. (C) Genomic abundance and gonadal expression level of TE superfamilies are positively correlated in both sexes.

### 2.3 Expression of TE-mapping piRNAs correlates with TE expression in the gonads

In both testes and ovaries, the length distribution of small RNAs includes a peak at 29 nucleotides (Figure 3A), and sequences up to 30 nt show a strong 5’-U bias at the first nucleotide position, consistent with expectations for the piRNA pool (Figure 3B). In ovaries, there is a second peak at 22 nucleotides. The relative expression of putative piRNAs (25–30 nt) is lower in females than in males. In contrast, the relative expression of ∼22 nt RNAs is higher in females than in males; 41%–59% of 22 nt RNAs correspond to known miRNAs in females, versus 37%–41% in males.

**Figure 3.**
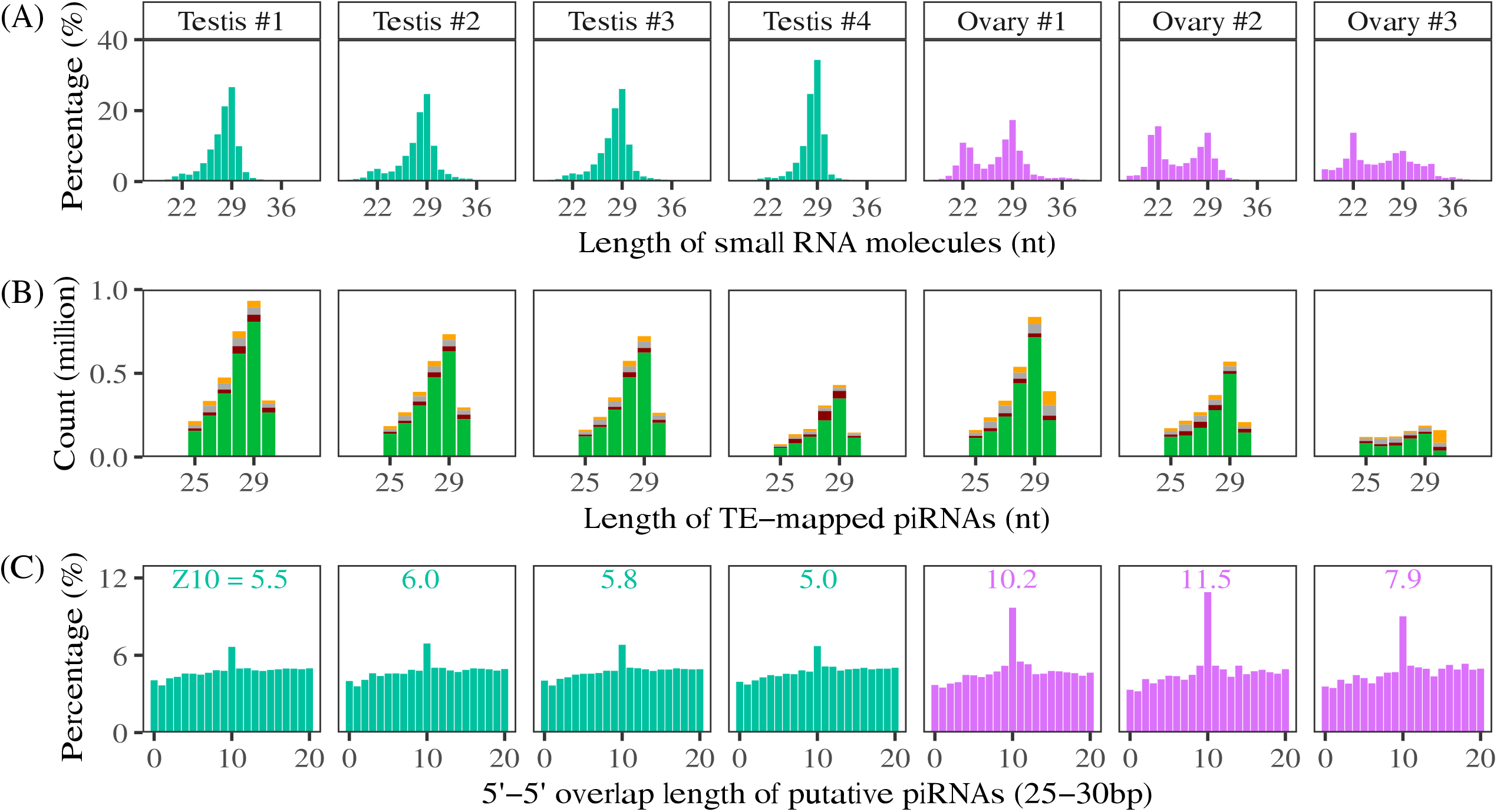
(A) The length distribution of small RNA molecules. (B) Composition of the first nucleotide (green means U) of small RNAs that could be mapped to TE transcripts in different genders. (C) The 5’ to 5’ overlapping of putative piRNA (25–30 bp) in gonads of males and females, a typical signature of ping-pong biogenesis. The number above the peak at the 10^th^ overlapping is the Z-score.

A higher percentage of total putative piRNAs map to TEs in females than in males; on average, 22.7% map in the antisense direction and 23.5% map in the sense direction in females, and 11.0% map in the antisense direction and 10.1% map in the sense direction in males. Considering unique putative piRNA sequences, 16.9% map in the antisense direction and 17.1% map in the sense direction in females, and 15.1% map in the antisense direction and 14.7% map in the sense direction in males. Overall, more total putative piRNAs map to TEs in males than females, although the ranges overlap (1,264,088–3,045,727 in males vs. 897,586–2,503,814 in females) (Figure 3B).

In both sexes, we identify a peak overlap length between TE-mapping sense and anti-sense piRNAs of 10 base pairs, consistent with ping-pong amplification of piRNAs in response to TE transcription (Figure 3C, Supplementary S5, S6). The strength of the ping-pong signal, indicated by the Z-scores of the 10-nt overlap, is greater in females.

piRNA expression is correlated with TE expression, measured at the TE superfamily level, in both females and males (Figure 4, Supplementary File S4). At higher levels of TE expression, females show a trend of having more piRNAs relative to TE expression level than males.

**Figure 4.**
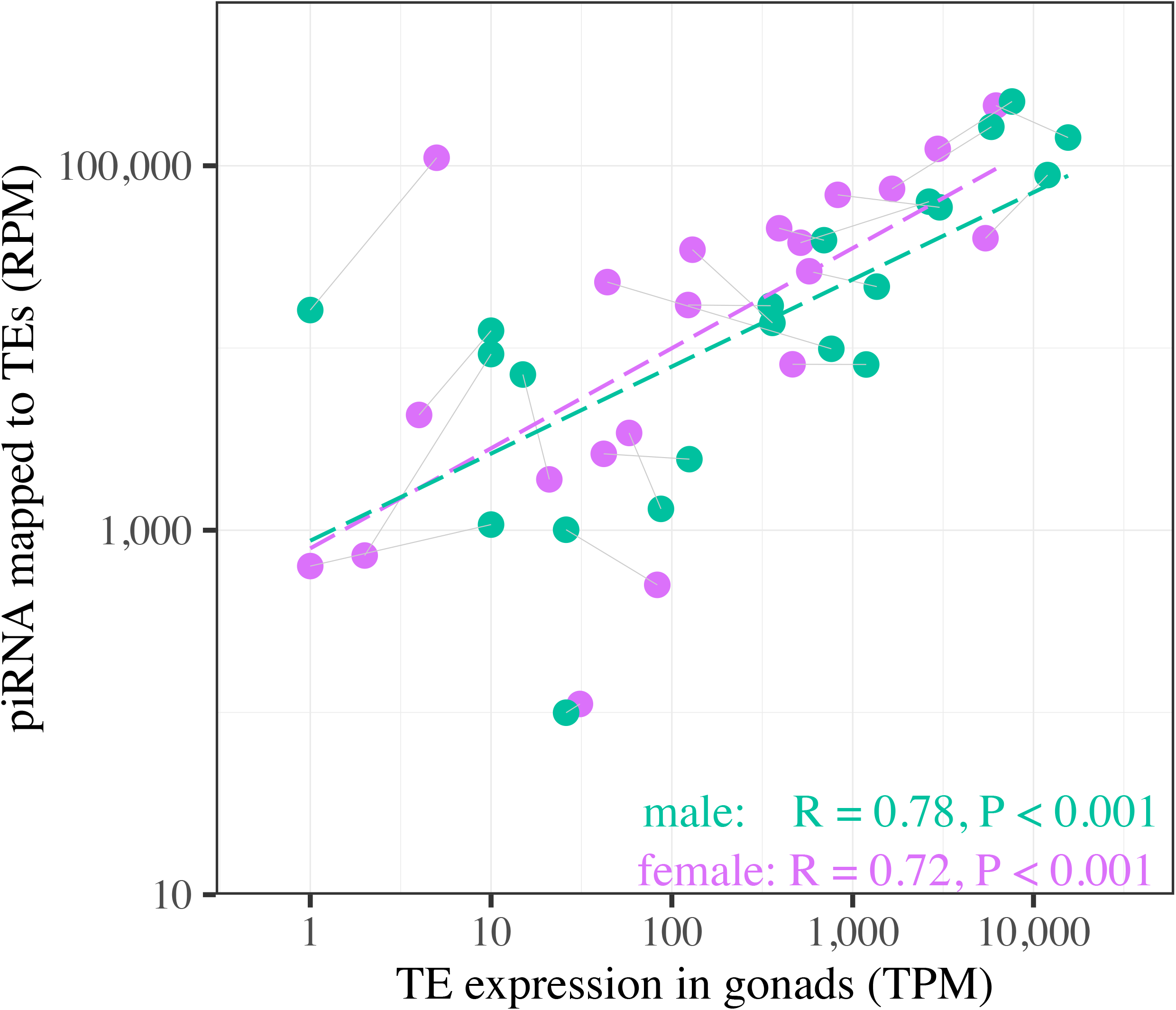
The relationship between TE expression and putative piRNAs (25–30 bp) mapped to TEs in both sexes. Gray lines connect male and female datapoints for the same TE superfamily.

### 2.4 Germline expression of piRNA pathway genes is higher in males in *R. sibiricus*, whereas KRAB-ZFP silencing and miRNA pathway genes are higher in females

The expression of piRNA pathway genes in *R. sibiricus* is higher in testes than in ovaries, measured both relative to miRNA pathway gene expression and as TPM (Figure 5; Supplementary Files 6,7). In contrast, the expression of genes establishing a repressive transcriptional environment (NuRD complex + related proteins) is comparable between the sexes relative to miRNA pathway gene expression and slightly higher in females measured as TPM. The expression of TRIM28 — which links KRAB-ZFP proteins to the NuRD complex + related proteins — is higher in females than males relative to miRNA expression, yet miRNA pathway expression levels (TPM) are slightly higher in females (consistent with higher miRNA expression, Figure 3; Supplementary Files 6,7). Taken together, these results suggest that males may rely more heavily on piRNA machinery to recruit repressive transcriptional machinery, whereas females may rely more heavily on KRAB-ZFP proteins to recruit repressive transcriptional machinery.

**Figure 5.**
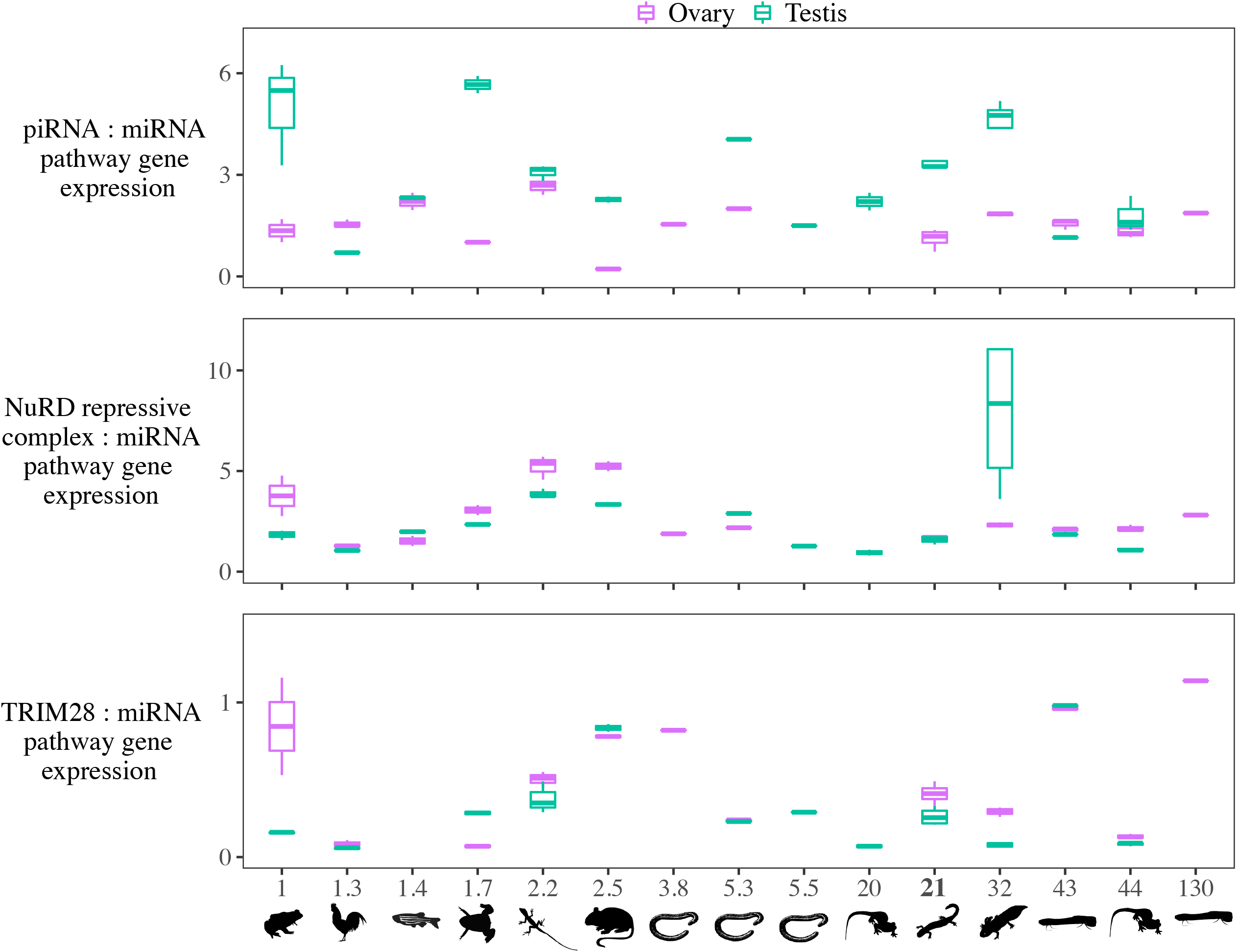
(A) The ratio of the summed expression of piRNA pathway genes, (B) NuRD and associated repressive complex genes, and (C) TRIM28 to the summed expression of miRNA pathway genes in species with diverse genome sizes. The species and their genome sizes (Gb) from left to right are: *Platyplectrum ornatum* (1), *Gallus gallus* (1.3), *Danio rerio* (1.4), *Xenopus tropicalis* (1.7), *Anolis carolinensis* (2.2), *Mus musculus* (2.5), *Geotrypetes seraphini* (3.8), *Rhinatrema bivittatum* (5.3), *Caecilia tentaculate* (5.5), *Pleurodeles waltl* (20), *Ranodon sibiricus* (21), *Ambystoma mexicanum* (32), *Protopterus annectens* (43), *Cynops orientalis* (44), and *Protopterus aethiiopicus* (∼130).

### 2.5 Relative expression of TE silencing pathways between males and females varies across species

Across species, higher piRNA pathway expression relative to miRNA pathway expression in males is seen in the majority of taxa across the range of genome sizes; the exceptions are *Gallus gallus* and *Protopterus annectens* (Figure 5, Supplementary File S6). Similarly, higher miRNA pathway expression in females (TPM) is seen in all taxa except *G. gallus* and *D. rerio*, although the difference is not always as pronounced as in *R. sibiricus* (Supplementary File S7). *Platyplectrum ornatus, Anolis carolinensis*, and *C. orientalis* show the same pattern of TE silencing expression differences between males and females as in *R. sibiricus*, with higher reliance on piRNA machinery in males and higher reliance on KRAB-ZFP in females. However, this pattern does not hold in other taxa (Figure 5).

### 2.6 Germline expression of TE silencing machinery does not correlate with genome size in either sex

Across the range of genome sizes from ∼1 Gb to ∼130 Gb, we find no correlations between genome size and 1) the expression of piRNA processing genes, 2) NuRD complex and associated genes establishing a repressive transcriptional environment, or 3) TRIM28. Interestingly, five of the six highest piRNA pathway expression levels are found in amphibians, the clade with the most variation in genome size (Figure 5, Supplementary File S9).

## 3 Discussion

### 3.1 Permissive TE environment despite comprehensive piRNA-mediated silencing

The transposable element community in *Ranodon sibiricus* is comparable in diversity to other gigantic amphibian genomes and shares many of the same abundant TE superfamilies (e.g., LTR/Gypsy, DIRS and LINE/L1) (Sun et al., 2012;Sun and Mueller, 2014;Wang et al., 2021a). However, *R. sibiricus* differs from other salamanders in having high levels of LINE/Jockey, a superfamily shown to be abundant in the caecilian *Ichthyophis bannanicus*, but rare in other salamanders (Wang et al., 2021a). Both genomic and transcriptomic data suggest that all TE superfamilies are potentially experiencing ongoing transposition in *R. sibiricus*. Thus, like other gigantic amphibian genomes, *R. sibiricus* appears to have attained its large size because of the expansion of multiple types of TEs, supporting the notion of amphibian genomes as permissive TE environments. At the same time, all of these putatively active TE superfamilies are targeted by piRNAs, and piRNA levels are correlated with TE expression at the superfamily level, consistent with patterns in *Drosophila* and suggesting that the scope, if not efficacy, of TE silencing in *Ranodon sibiricus* is comparable to other species (Kelleher and Barbash, 2013;Saint-Leandre et al., 2020).

### 3.2 Sex-biased TE expression and silencing

Transposable element expression is higher in *R. sibiricus* testes than ovaries, a pattern also reported in *Drosophila* (Wei et al., 2022) and *Oryzias latipes* (the medaka fish) (Saint-Leandre et al., 2020;Dechaud et al., 2021), but opposite the pattern reported in *Corvus corone* (carrion crow) (Warmuth et al., 2022) and different from the non-sex-biased TE expression in the fire-bellied newt *Cynops orientalis* (Carducci et al., 2021). In *Drosophila*, testes expression levels of TE-mapping piRNAs and piRNA pathway genes, as well as the ping-pong signature, are lower than ovary expression levels, suggesting that lower male piRNA-mediated silencing contributes to higher male TE expression (Saint-Leandre et al., 2020;Chen et al., 2021). In *O. latipes*, on the other hand, testes expression levels of TE-mapping piRNAs are higher than ovary expression levels — a pattern also observed in zebrafish — suggesting that higher male piRNA-mediated silencing may actually be correlated with high male TE expression in these two fish species (Houwing et al., 2007;Kneitz et al., 2016). In *R. sibiricus*, TE-mapping piRNA counts are similar between the sexes, despite higher relative expression of putative piRNAs in males than females (Figure 3a,b). However, the ping-pong amplification signature is higher in females, as is the number of piRNAs per expressed TE transcript, suggestive of a more robust piRNA-directed silencing response in females (Figure 3c, 4). On the other hand, the piRNA pathway protein expression levels are higher in males, suggesting the opposite case of a more robust response in males (Figure 5).

Sex-specific differences in gonadal TE expression have also been explained by factors other than variation in TE silencing; for example, in systems with heteromorphic sex chromosomes, sex-biased TE expression has been attributed to different TE dynamics on the sex-limited chromosome (Y in XY systems, or W in ZW systems). TE abundance is higher on sex-limited chromosomes because of lower effective population size, lack of recombination, and lower gene density. In addition, TE expression per locus has been shown to be higher on the sex-limited chromosome itself — as well as genome-wide — in the heterogametic sex in *Drosophila* (Y, males) and in crows (*Corvus corone*; W, females) (Wei et al., 2020;Warmuth et al., 2022). The mechanism for higher TE expression in the heterogametic sex in these cases remains incompletely understood, but may involve TEs affecting their own genome-wide regulation *in trans* or the heightened conflict between creating a repressive chromatin state to silence TEs while maintaining open chromatin to allow genic transcription on a degenerating chromosome (Wei et al., 2020;Warmuth et al., 2022). Neither *Ranodon sibiricus* nor *Oryzias latipes* has heteromorphic sex chromosomes (Hillis and Green, 1990;Matsuda et al., 2002;Evans et al., 2012;Perkins et al., 2019), yet both show sex-biased TE expression, and only *Oryzias* shows unambiguous sex-biased TE-targeting piRNA expression; taken together, these data reveal that the difference in TE expression between sexes reflects different underlying causes across species. The relationships among sex-biased TE expression, sex determination, and TE silencing are an important target for future research, but irrespective of the underlying mechanisms, the sex with higher TE activity contributes more to the species’ genomic TE load. In *R. sibiricus*, that sex is the male, provided that TE expression is a reasonable proxy for transposition.

### 3.3 The TE-silencing arms race dynamic across genome sizes

Transposable elements and the silencing machinery of their hosts are engaged in an arms race in which a novel TE family initially proliferates, the host evolves silencing based on TE sequence identification, and the TE subsequently diverges to evade silencing — or that particular TE remains permanently silenced, but a novel TE invades the host genome and begins the cycle anew (Luo et al., 2020;Zhang et al., 2020;Said et al., 2022;Wei et al., 2022). If balanced by deletion of TE sequences, this arms-race dynamic can be associated with fairly stable genome size over evolutionary timescales, despite turnover in TE content (Kapusta et al., 2017). To yield an overall evolutionary trend in genome size, the long-term balance between TE insertion and deletion has to become skewed in favor of one or the other, with deletion bias leading to genome contraction and insertion bias leading to genome expansion (Nam and Ellegren, 2012).

It has been suggested in the literature that large genomes may be manifestations of an arms race between TEs and the silencing of their hosts, but that this arms race involves the TE community as a whole rather than individual TE families — and the species with the gigantic genome has been interpreted as both the current “attacker” and “defender” in the arms race. For example, Meyer et al (2021) suggested that TE silencing machinery did not adapt to reduce TE expansion in the Australian lungfish, based on high genomic TE abundance and ongoing TE expression. In the strawberry poison dart frog, which has a moderately expanded genome size of 6.76 Gb, a widespread failure to silence TEs was suggested based on high genomic TE abundance, germline TE expression of diverse TEs, and the presence of an identified *piwi2* and *piwi3* transcript in only one individual sampled each (Rogers et al., 2018). In the salamander *Desmognathus fuscus*, less comprehensive piRNA pathway-mediated TE silencing was suggested based on a relatively low percentage of TE-mapping piRNAs (Madison-Villar et al., 2016). In all three of these examples, the TEs are suggested to be currently on the attack. On the other hand, in the African lungfish, Wang et al. (2021) suggested that the KRAB-ZFP TE transcriptional silencing machinery has expanded in scope in response to the high genomic TE load. Similarly, in the fire-bellied newt *Cynops orientalis*, it was suggested that TE silencing is now enhanced in response to high TE load, which accumulated in the past during a period of increased TE mobilization (Carducci et al., 2021). In both of these examples, the TEs are suggested to be currently on the defensive. Thus, two opposite predictions for TE silencing in gigantic genomes — either increased or decreased — have been proposed to be met in different extant organisms, albeit with datasets that are not necessarily comparable. This apparent conflict can be resolved by considering that the attack/defense status of TEs and their silencing reflect where the lineage currently exists in the dynamic cycle between TE and host dominance. Our results revealing no consistent pattern in TE silencing pathway expression levels and genome size (Figure 5) are consistent with this interpretation. At large sizes, it is not the size of the genome itself that likely predicts the efficacy of the TE silencing machinery, but more likely the directional trend in genome size evolution; genomes that are contracting are more likely to have effective TE silencing, whereas genomes undergoing expansion are more likely to have reduced TE silencing. In the absence of comparable data (including small RNA, TE expression and amplification, and silencing pathway expression) for other species with known trends in genome size evolution, we opt for a conservative position and do not infer *Ranodon sibiricus’* current position in the TE/host dynamic arms race cycle.

The mechanisms by which global TE silencing mechanisms can be subverted by a community of TEs, and then evolve to regain stricter control, are not yet well-understood. A few studies in invertebrates have begun to reveal differences in TE and silencing dynamics in genomes of different sizes. In a pairwise comparison of grasshopper species with different genome sizes, Liu et al. revealed that the species with the larger genome had higher TE expression, lower piRNA abundance, and lower expression of the piRNA biogenesis gene *HENMT*, which suggested that lower piRNA-mediated TE silencing was permissive to higher TE activity and genome expansion (Liu et al., 2022). A comparison between *Drosophila melanogaster* and the mosquito *Aedes aegypti*, which has a larger genome (1.38 Gb versus 180 Mb), revealed that the mosquito has a higher TE load and a smaller percentage of TE-mapping piRNAs (Arensburger et al., 2011). Future studies that leverage the large range of genome sizes present in vertebrates, emphasizing comprehensive across-species data on TE activity and TE silencing in a phylogenetic context to allow ancestral genome size reconstruction, will continue to shed light on how TEs and their hosts coevolve to achieve gigantic genomes.

## 4 Materials and methods

### 4.1 Specimen information

We collected three male adult *Ranodon sibiricus* from the wild of Wenquan County, Xinjiang Uygur Autonomous Region of China, and egg-hatched and raised one male and four females in an aquarium of Xinjiang Normal University from eggs originally collected from the same field site as the wild males. All these individuals were collected during the breeding season of August, 2017, and all adults had a snout-tail length of 16–21 cm and a body mass of 12–35 g prior to euthanasia (Supplementary File S1). Wild-caught adults were euthanized upon return to the laboratory and were not kept alive in captivity. Collection, hatching, and euthanasia were performed following Animal Care & Use Protocols approved by Chengdu Institute of Biology, Chinese Academy of Sciences.

### 4.2 Genome size estimation

Blood smears were prepared from a formalin-fixed specimen of *Ranodon sibiricus* and nuclear area was measured from Fuelgen-stained red blood cell nuclei using the ImagePro® image analysis program (Itgen et al., 2022). Blood smears of the reference standards *Ambystoma mexicanum* (34 Gb; Gregory, 2022) and the Iberian ribbed newt *Pleurodeles waltl* (20 Gb; Gregory 2022) were prepared and analyzed at the same time under the same conditions.

### 4.3 Genomic shotgun library creation, sequencing, and assembly

Total DNA was extracted from muscle tissue using the modified low-salt CTAB extraction of high-quality DNA procedure (Arseneau et al., 2017). DNA quality and concentration were assessed using agarose gel electrophoresis and a NanoDrop Spectrophotometer (ThermoFisher Scientific, Waltham, MA), and a PCR-free library was prepared using the NEBNext Ultra DNA Library Prep Kit for Illumina. Sequencing was performed on one lane of a Hiseq 2500 platform (PE250). Library preparation and sequencing were performed by the Beijing Novogene Bioinformatics Technology Co. Ltd. Raw reads were quality-filtered and adaptor-trimmed using Trimmomatic-0.39 (Bolger et al., 2014) with default parameters. In total, the genomic shotgun dataset included 11,960,858 reads. After filtering and trimming, 11,168,678 reads covering a total length of 2,314,096,923 bp remained. Thus, the sequencing coverage is about 10.1% (0.1X coverage). Filtered, trimmed reads were assembled into contigs using dipSPAdes 3.12.0 (Bankevich et al., 2012) with default parameters, yielding 478,991 contigs with an N50 of 447 bp and a total length of 249,425,929 bp.

### 4.4 Mining and classification of repeat elements

The PiRATE pipeline was used as in the original publication (Berthelier et al., 2018), including the following steps: **1)** Contigs representing repetitive sequences were identified from the assembled contigs using similarity-based, structure-based, and repetitiveness-based approaches. The similarity-based detection programs included RepeatMasker v-4.1.0 (http://repeatmasker.org/RepeatMasker/, using Repbase20.05_REPET.embl.tar.gz as the library instead) and TE-HMMER (Eddy, 2011). The structural-based detection programs included LTRharvest (Ellinghaus et al., 2008), MGEScan non-LTR (Rho and Tang, 2009), HelSearch (Yang et al., 2009), MITE-Hunter (Han and Wessler, 2010), and SINE-finder (Wenke et al., 2011). The repetitiveness-based detection programs included TEdenovo (Flutre et al., 2011) and RepeatScout (Price et al., 2005). **2)** Repeat consensus sequences (*e*.*g*., representing multiple subfamilies within a TE family) were also identified from the cleaned, filtered, and unassembled reads with dnaPipeTE (Goubert et al., 2015) and RepeatModeler (http://www.repeatmasker.org/RepeatModeler/). **3)** Contigs identified by each individual program in steps 1 and 2, above, were filtered to remove those < 100 bp in length and clustered with CD-HIT-est (Li and Godzik, 2006) to reduce redundancy (100% sequence identity cutoff). This yielded a total of 155,999 contigs. **4)** All 155,999 contigs were then clustered together with CD-HIT-est (100% sequence identity cutoff), retaining the longest contig and recording the program that classified it. 46,090 contigs were filtered out at this step. **5**) The remaining 109,909 repeat contigs were annotated as TEs to the levels of order and superfamily in Wicker’s hierarchical classification system (Wicker et al., 2007), modified to include several recently discovered TE superfamilies using PASTEC (Hoede et al., 2014), and checked manually to filter chimeric contigs and those annotated with conflicting evidence (Supplementary File S2). **6**) All classified repeats (“known TEs” hereafter), along with the unclassified repeats (“unknown repeats” hereafter) and putative multi-copy host genes, were combined to produce a *Ranodon*-derived repeat library. **7**) For each superfamily, we collapsed the contigs to 95% and 80% sequence identity using CD-HIT-est to provide an overall view of within-superfamily diversity; 80% is the sequence identity threshold used to define TE families (Wicker et al., 2007).

### 4.5 Characterization of the overall repeat element landscape

Overlapping paired-end shotgun reads were merged using PEAR v.0.9.11 (Zhang et al., 2014) with the following parameter values based on our library insert size and trimming parameters: min-assemble-length 36, max-assemble-length 490, min-overlap size 10. After merging, 7,385,166 reads remained (including both merged and singletons), with an N50 of 388 bp and total length of 1,997,175,501 bp. To calculate the percentage of the *R. sibiricus* genome composed of different TEs, these shotgun reads were masked with RepeatMasker v-4.1.0 using two versions of our *Ranodon*-derived repeat library: one that included the unknown repeats and the other that excluded them. In both cases, simple repeats were identified using the Tandem Repeat Finder module implemented in RepeatMasker. The overall results were summarized at the levels of TE class, order, and superfamily.

### 4.6 Measuring diversity of the genomic TE community

Unknown repeats were excluded from the analysis, as were TEs that could only be annotated down to the level of Class. Simpson’s diversity index is expressed as the variable D, calculated by: 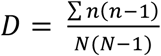 (Simpson, 1949). D is the probability that two individuals at random pulled from a community will be from the same species. We report 1 – D, or the Gini-Simpson’s index, which is more intuitive. The Shannon’s diversity index H is calculated by: 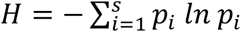 (Shannon, 1948). The higher the value of H, the greater the diversity.

### 4.7 Amplification history of TE superfamilies

To summarize the overall amplification history of TE superfamilies and test for ongoing activity, the perl script parseRM.pl (Kapusta et al., 2017) was used to parse the raw output files from RepeatMasker (.align) and report the sequence divergence between each read and its respective consensus sequence (parameter values = -l 50,1 and -a 5). The repeat library used to mask the reads comprised the 55,327 TE contigs classified by the PiRATE pipeline and clustered at 100% sequence identity. Each TE superfamily is therefore represented by multiple consensus sequences corresponding to the family and subfamily TE taxonomic levels (*i*.*e*., not the distant common ancestor of the entire superfamily). For each superfamily, histograms were plotted to summarize the percent divergence of all reads from their closest (*i*.*e*., least divergent) consensus sequence. These histograms do not allow the delineation between different amplification dynamics scenarios (*i*.*e*., a single family with continuous activity *versus* multiple families with successive bursts of activity). Rather, these global overviews were examined for overall shapes consistent with ongoing activity (*i*.*e*., the presence of TE loci < 1% diverged from the ancestral sequence and a unimodal, right-skewed, J-shaped, or monotonically decreasing distribution).

### 4.8 Transcriptome library creation, sequencing, assembly, and TE annotation

Total RNA was extracted separately from testes (*n* = 4) and ovary (*n* = 4) tissues using TRIzol (Invitrogen). For each sample, RNA quality and concentration were assessed using agarose gel electrophoresis, a NanoPhotometer spectrophotometer (Implen, CA), a Qubit 2.0 Fluorometer (ThermoFisher Scientific), and an Agilent BioAnalyzer 2100 system (Agilent Technologies, CA), requiring an RNA integrity number (RIN) of 8.5 or higher; one ovary sample failed to meet these quality standards and was excluded from downstream analyses. Sequencing libraries were generated using the NEBNext Ultra RNA Library Prep Kit for Illumina following the manufacturer’s protocol. After cluster generation of the index-coded samples, the library was sequenced on one lane of an Illumina Hiseq 4000 platform (PE 150). Transcriptome sequences were filtered using Trimmomatic-0.39 with default parameters (Bolger et al., 2014). 30,848,170 to 39,695,323 reads were retained for each testis or ovary sample, and in total, 290,925,984 reads remained, with a total length of 42,385,060,050 bp. Remaining reads of all testes and ovary samples were combined and assembled using Trinity 2.12.0 (Haas et al., 2013), yielding 573,144 contigs (i.e., putative assembled transcripts). Contigs were clustered using CD-hit-est (95% identity). Completeness of this final *de novo* transcriptome assembly were assessed using the BUSCO pipeline (Simao et al., 2015).

Expression levels of contigs in each sample were measured with Salmon (Patro et al., 2017), and contigs with no raw counts were removed. To annotate the remaining contigs containing autonomous TEs, BLASTp and BLASTx were used against the Repbase Database (downloaded on January 5, 2022) with an E-value cutoff of 1E-5 and 1E-10, respectively. The aligned length coverage was set to exceed 80% of the queried transcriptome contigs. To annotate contigs containing non-autonomous TEs, RepeatMasker was used with our *Ranodon*-derived genomic repeat library of non-autonomous TEs (LARD-, TRIM-, MITE-, and SINE-annotated contigs) and the requirement that the transcriptome/genomic contig overlap was > 80 bp long, > 80% identical in sequence, and covered > 80% of the length of the genomic contig. Contigs annotated as conflicting autonomous and non-autonomous TEs were filtered out.

To identify contigs that contained endogenous *R. sibiricus* genes, the Trinotate annotation suite (Bryant et al., 2017) was used with an E-value cutoff of 1E-5 for both BLASTx and BLASTp against the Uniport database, and 1E-5 for HMMER against the Pfam database (Wheeler and Eddy, 2013). To identify contigs that contained both a TE and an endogenous gene (i.e., putative cases where a TE and a gene were co-transcribed on a single transcript), all contigs that were annotated both by Repbase and Trinotate were examined, and the ones annotated by Trinotate to contain a TE-encoded protein (*i*.*e*., the contigs where Repbase and Trinotate annotations were in agreement) were not further considered. The remaining contigs annotated by Trinotate to contain a non-TE gene (*i*.*e*., an endogenous *Ranodon* gene) and also annotated either by Repbase to include a TE-encoded protein or by RepeatMasker to include a non-autonomous TE were identified for further examination and expression-based analysis.

### 4.9 Germline TE expression quantification in males and females

Expression levels of the individual TE superfamilies were calculated by averaging the TPM values among replicates of each sex and then summing the average TPM of all contigs annotated to each superfamily. For TE superfamilies detected in both the genomic and transcriptomic datasets, we tested for a relationship between genomic abundance and expression levels in each sex using linear regression on log-transformed data.

To identify differentially expressed contigs between males and females, DESeq2 (Love et al., 2014) was used with an adjusted P-value cut off of 0.05. Among the 15,011 total differentially expressed transcripts between sexes (including TEs, endogenous genes, and unannotated contigs), 869 were TEs, representing 18 superfamilies and other unknown TEs. Superfamilies with fewer than 10 differentially expressed transcripts between sexes were removed, leaving 9 superfamilies; for each, we tested for a difference in expression between males and females using a t-test.

### 4.10 Identification of putative piRNAs from small RNA-seq data

Small RNA libraries were prepared for each sample using the NEBNext® Multiplex Small RNA Library Prep Set for Illumina® (NEB, USA) following the manufacturer’s recommendations, and index codes were added to attribute sequences to each sample. Briefly, the NEB 3’ SR Adaptor (5’-AGATCGGAAGAGCACACGTCT-3’) was ligated to the 3’ end of small RNA molecules. After the 3’ ligation reaction, the SR RT Primer was hybridized to the excess 3’ SR Adaptor (that remained free after the 3’ ligation reaction), transforming the single-stranded DNA adaptor into a double-stranded DNA molecule (dsDNAs). This step was important for preventing adaptor-dimer formation, and because dsDNAs are not substrates for ligation mediated by T4 RNA Ligase 1, they therefore would not ligate to the 5’ SR Adaptor (5’-GTTCAGAGTTCTACAGTCCGACGATC-3’) in the subsequent ligation step. The 5’ end adapter was then ligated to the 5’ ends of small RNA molecules. First strand cDNA was synthesized using M-MuLV Reverse Transcriptase (Rnase H). PCR amplification was performed using LongAmp Taq 2X Master Mix, SR Primer for Illumina, and index primers. PCR products were purified on an 8% polyacrylamide gel (100V, 80 min). DNA fragments corresponding to ∼140-160 bp (the length of a small noncoding RNA plus the 3’ and 5’ adaptors) were recovered and dissolved in 8 μL elution buffer. Library quality was assessed on the Agilent Bioanalyzer 2100 using DNA High Sensitivity Chips. Clustering of index-coded samples was performed on a cBot Cluster Generation System using the TruSeq SR Cluster Kit v3-cBot-HS (Illumina) according to the manufacturer’s instructions. After cluster generation, libraries were sequenced on an Illumina Hiseq 2500 platform (SE50).

We filtered low-quality sequences using the fastq_quality_filter (-q 20, -p 90) in the FASTX-Toolkit v0.0.13 (http://hannonlab.cshl.edu/fastx_toolkit/). We removed the adapter sequence with a minimum overlap of 10 bp from the 3’-end, discarded untrimmed reads, and selected those with a minimum length of 18 bp and a maximum length of 40 bp (after cutting adapters) and no Ns using cutadapt v2.8 (Martin, 2011). Reads mapping to the mitochondrial genome (NCBI code: AJ419960) and riboRNAs (NCBI codes: DQ283664, AJ279506, MH806872) were identified and filtered out using Bowtie v1.1.0 (Langmead et al., 2009). Overall, more reads were filtered out based on length and rRNA identity in females than males (Supplementary File S5). miRNAs 21-24 nt in length were annotated using Bowtie v1.1.0 to identify hits to miRbase 22.1 (Kozomara et al., 2019) in each male and female.

To test for the predicted bias towards U at the first 5’ nucleotide position of piRNAs, we calculated the proportion of small RNAs with each nucleotide in the first position. Based on this result, and the overall length distribution of RNAs between 18 and 40 nt, we conservatively defined putative piRNAs as those ranging from 25-30 nt, and we selected these using the seqkit software (Shen et al., 2016). We mapped these putative piRNAs to our transcriptome assembly using Bowtie and identified piRNAs that map to autonomous TEs (i.e. those that include transposition-related ORFs) in the sense and antisense orientations.

### 4.11 Ping-pong signature analysis

Secondary piRNA biogenesis associated with piRNA-targeted post-transcriptional TE silencing produces a distinctive “ping-pong signature” in the piRNA pool, which consists of a 10 bp overlap between the 5’ ends of antisense and sense piRNAs. The ping-pong signature for each individual was analyzed using the following approach: First, TE transcripts that were not mapped by both sense-oriented and antisense-oriented piRNAs were filtered out using Bowtie, allowing 0 mismatches for sense mapping under the assumption that piRNAs derived directly from an RNA target should have the identical sequence (Teefy et al., 2020) and 3 mismatches for antisense mapping because cleavage of RNA targets can occur with imperfect base-pairing (Zhang et al., 2015). Second, the fractions of overlapping pairs of sense/antisense piRNAs corresponding to specific lengths, as well as the Z-score measuring the significance of each ping-pong signature, were generated using the script of 1_piRNA_and_Degradome_Counts.RMD (Teefy et al., 2020).

### 4.12 Putative piRNAs targeting TE superfamilies

To estimate levels of piRNAs targeting each TE superfamily, putative piRNAs were mapped to the TE transcripts using the ‘align_and_estimate_abundance.pl’ script of Trinity (Haas et al., 2013). Reads per million (RPM) values were calculated for each TE contig and then averaged across individuals of each sex. For each sex, overall putative piRNA levels targeting each TE superfamily were calculated by summing across all contigs annotated to the same TE superfamily. We tested for a correlation between TE superfamily expression level and targeting piRNA abundance using linear regression on the log-transformed variables.

### 4.13 Germline TE silencing pathway expression across genome sizes

To test for an association between overall piRNA or KRAB-ZFP pathway activity and genome size, we first compiled male and female gonad RNA-Seq datasets for vertebrates of diverse genome sizes, including *Platyplectrum ornatum* (ornate burrowing frog), *Gallus gallus* (chicken), *Danio rerio* (zebrafish), *Xenopus tropicalis* (Western clawed frog), *Anolis carolinensis* (green anole), *Mus musculus* (mouse), *Geotrypetes seraphini* (Gaboon caecilian), *Rhinatrema bivittatum* (two-lined caecilian), and *Caecilia tentaculata* (bearded caecilian) spanning genomes sizes from 1.0 – 5.5 Gb, and *Pleurodeles waltl* (the Iberian ribbed newt), *Ambystoma mexicanum* (the Mexican axolotl), *Cynops orientalis* (the fire-bellied newt), *Protopterus annectens*, and *P. aethiopicus* (African and marbled lungfishes) spanning genome sizes from 20 – ∼130 Gb (Supplementary File S8). We performed de novo assemblies using the same pipeline as for *R. sibiricus* on all obtained datasets.

We identified transcripts of 21 genes receiving a direct annotation of piRNA processing in vertebrates in the Gene Ontology knowledgebase that were present in the majority of our target species: ASZ1, BTBD18 (BTBDI), DDX4, EXD1, FKBP6, GPAT2, HENMT1 (HENMT), MAEL, MOV10l1 (M10L1), PIWIL1, PIWIL2, PIWIL4, PLD6, TDRD1, TDRD5, TDRD6, TDRD7, TDRD9, TDRD12 (TDR12), TDRD15 (TDR15), and TDRKH. In addition, we identified transcripts of 14 genes encoding proteins that create a transcriptionally repressive chromatin environment in response to recruitment by PIWI proteins or KRAB-ZFP proteins, 12 of which received a direct annotation of NuRD complex in the Gene Ontology knowledgebase and 2 of which were taken from the literature: CBX5, CHD3, CHD4, CSNK2A1 (CSK21), DNMT1, GATAD2A (P66A), MBD3, MTA1, MTA2, RBBP4, RBBP7, SALL1, SETDB1 (SETB1), and ZBTB7A (ZBT7A) (cite Ecco et al 2017) (Wang et al., 2022). Finally, we identified TRIM28, which bridges this repressive complex to TE-bound KRAB-ZFP proteins in lobe-finned fishes (Ecco et al., 2017). For comparison, we identified transcripts of 14 protein-coding genes receiving a direct annotation of miRNA processing in vertebrates in the Gene Ontology knowledgebase, which we did not predict to differ in expression based on genome size: ADAR (DSRAD), AGO1, AGO2, AGO3, AGO4, DICER1, NUP155 (NU155), PUM1, PUM2, SNIP1, SPOUT1 (CI114), TARBP2 (TRBP2), TRIM71 (LIN41), and ZC3H7B. Expression levels for each transcript in each individual were measured with Salmon (Patro et al., 2017) (Supplementary File S9).

As a proxy for overall piRNA silencing activity, for each individual, we calculated the ratio of total piRNA pathway expression (summed TPM of 21 genes) to total miRNA pathway expression (summed TPM of 14 genes). As a proxy for transcriptional repression driven by both the piRNA pathway and KRAB-ZFP binding activity, we calculated the ratio of total transcriptional repression machinery expression (summed TPM of 14 genes) to total miRNA pathway expression. Finally, we calculated the ratio of TRIM28 expression to total miRNA pathway expression for each individual. We also calculated these ratios with a more conservative dataset allowing for no missing genes; this yielded 15 piRNA pathway genes, 9 KRAB-ZFP genes, and 13 miRNA genes. We plotted these ratios to reveal any relationship between TE silencing pathway expression and genome size.

## Supporting information

Supplemental Files 1 through 8

## 5 Data availability statement

Genomic shotgun and transcriptome sequences have been deposited in the Genome Sequence Archive at the National Genomics Data Center, Beijing Institute of Genomics, Chinese Academy of Sciences/China National Center for Bioinformation (GSA: CRA008892, CRA008899, CRA008900), and are publicly accessible at http://bigd.big.ac.cn/gsa.

## 6 Author contributions

**Jie Wang**: Conceptualization, Methodology, Software, Formal analysis, Investigation, Resources, Data curation, Writing - original draft, Writing - review & editing, Visualization, Funding acquisition. **Liang Yuan**: Specimen collection. **Jiaxing Tang Jiongyu Liu, Cheng Sun, Michael W. Itgen, & Guiying Chen**: Methodology, Software. **Stanley K. Sessions**: Investigation, Resources. **Guangpu Zhang**: Methodology, Formal analysis, Writing - original draft, Writing - review & editing, Visualization. **Rachel Lockridge Mueller**: Conceptualization, Formal analysis, Investigation, Resources, Writing - original draft, Writing - review & editing, Supervision, Project administration, Funding acquisition. All authors read and approved the final manuscript.

## 8 Acknowledgements

We gratefully acknowledge Ye Xu and Yuzhou Gong for target tissue dissection; Xiuling Wang for help in field work; Jianping Jiang, Ava Louise Haley, and Chaochao Yan for their expertise in facilitating data analysis.

