## Supplemental Files 1 through 8 for "Transposable element and host silencing activity in gigantic genomes"

### S1 Samples and data used in this experiment

| Tissue | Code | Sex | Mass: g | STL: cm | Collection date | Source | Genomic sequencing | | RNA quality | | | | mRNA | | small RNA | |
|---|---|---|---|---|---|---|---|---|---|---|---|---|---|---|---|---|
| | | | | | | | Raw reads | Accession code[1] | OD260/280 | OD260/230 | 28S/18S | RIN | Raw reads | Accession code[2] | Raw reads | Accession code[3] |
| Testis#1 | WQ07 | ♂ | 16.83 | 17.252 | 2017.8.22. | wild | | | 1.77 | 2.75 | 1.7 | 9.6 | 37,888,360 | CRR610084 | 19,717,749 | CRR610091 |
| Testis#2 | WQ08 | ♂ | 19.79 | 18.192 | 2017.8.22. | wild | | | 1.73 | 0.95 | 1.1 | 8.9 | 35,386,396 | CRR610085 | 21,815,001 | CRR610092 |
| Testis#3 | WQ10 | ♂ | 12.02 | 16.441 | 2017.8.22. | wild | | | 2.00 | 3.22 | 1.7 | 9.4 | 34,854,594 | CRR610086 | 16,177,680 | CRR610093 |
| Testis#4 | WQ05 | ♂ | 20.48 | 15.591 | 2017.8.22. | captive | | | 1.70 | 2.13 | 1.7 | 9.3 | 31,732,246 | CRR610083 | 19,005,099 | CRR610090 |
| Ovary#1 | WQ03 | ♀ | 32.85 | 21.001 | 2017.8.22. | captive | 11,960,858 | CRR609958 | 1.96 | 0.68 | 1.1 | 9.3 | 38,466,202 | CRR610080 | 17,965,762 | CRR610088 |
| Ovary#2 | WQ02 | ♀ | 28.61 | 20.804 | 2017.8.22. | captive | | | 1.64 | 0.49 | 1.5 | 9.6 | 40,660,732 | CRR610079 | 15,836,634 | CRR610087 |
| Ovary#3 | WQ04 | ♀ | 35.37 | 20.437 | 2017.8.22. | captive | | | 1.64 | 0.22 | 0.8 | 8.7 | 39,418,784 | CRR610081 | 14,063,357 | CRR610089 |
| Ovary#4 | WQ06 | ♀ | 23.57 | 18.149 | 2017.8.22. | captive | | | 1.66 | 0.15 | 0.1 | 7.7[4] | 39,229,748 | CRR610082 | | |

[1]https://ngdc.cncb.ac.cn/gsa/s/5d3dHx5W

[2]https://ngdc.cncb.ac.cn/gsa/s/p15v402F

[3]https://ngdc.cncb.ac.cn/gsa/s/X3i83677

[4]This individuals was deleted from the small RNA analysis because of its low RIN value.

### S2 Types of genomic repeats annotated by PASTEC

| Type | No. of Contigs |
|---|---|
| Potential multiple host gene | 275 |
| SSR | 2,448 |
| Potential Chimeric | 1,088 |
| Conflicting evidence | 20 |
| noCat (unknown repeats) | 51,857 |
| known TEs | 54,221 |
| in total | 109,909 |

### S3 Expression levels of genes and TEs (left, females; right, males)

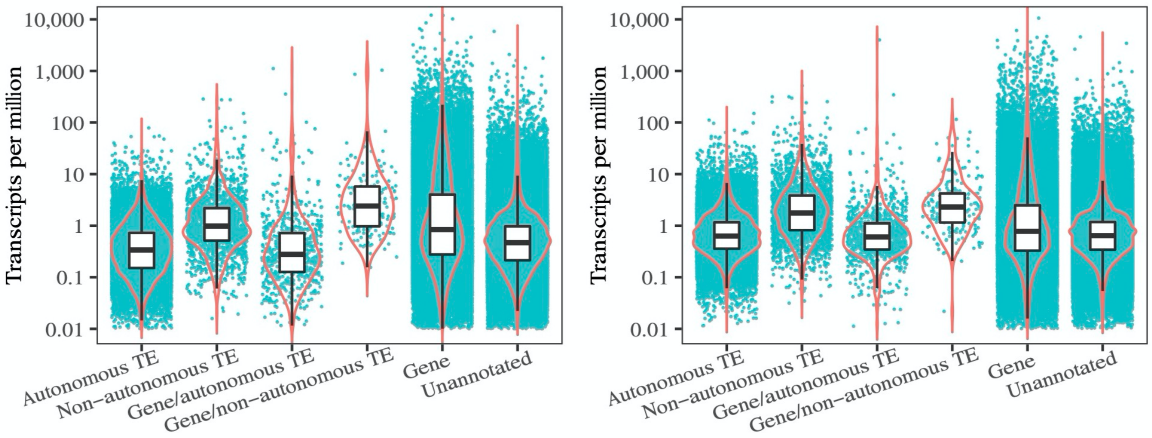

### S4 Genomic content, expression level of TEs, and putative piRNAs mapping to TEs superfamilies

| Superfamily | TE Content in genome: % | TE expression | | Putative piRNAs mapping to TEs | |
|---|---|---|---|---|---|
| | | females (TPM) | males (TPM) | female (RPM) | males (RPM) |
| LTR/Gypsy | 3.85 | 826.9 | 3024.4 | 69264.4 | 59,160.3 |
| LTR/ERV | 0.41 | 391.9 | 693.9 | 45,392.8 | 39,045.4 |
| LTR/Copia | 0.1 | 30.9 | 25.7 | 111.3 | 99.7 |
| LTR/Retrovirus | | 0.7 | 9.9 | 635.8 | 1,074 |
| DIRS/DIRS | 4.44 | 5427 | 11961.7 | 40,038.7 | 88,936.7 |
| PLE/Penelope | 0.09 | 122.8 | 351.6 | 17,211.8 | 17,062.2 |
| LINE/Jockey | 9.69 | 6206.5 | 15535.2 | 214,022.6 | 142,910.1 |
| LINE/L1 | 5.04 | 2954.2 | 7609.5 | 123,807.1 | 225,250.5 |
| LINE/I | 0.09 | 41.5 | 124.8 | 2,625.7 | 2,454.6 |
| LINE/RTE | 0.12 | 130 | 360.1 | 34,633.8 | 13,732 |
| SINE/5S | 0.23 | 5.3 | 1.3 | 110,673.1 | 16,152.2 |
| TRIM | 3.8 | 1650.8 | 5846.7 | 74,725.9 | 163,669.3 |
| LARD | 0.15 | 515.1 | 2643.3 | 37,826.3 | 63,559.3 |
| TIR/PIF-Harbinger | 2.98 | 575.2 | 1358.8 | 26,374.9 | 21,717.9 |
| TIR/hAT | 1.15 | 83.3 | 26.3 | 501.5 | 1,006.6 |
| TIR/Tc1-Mariner | 0.18 | 57.9 | 87.2 | 3,417.5 | 1,310.2 |
| TIR/PiggyBac | 0.05 | 21.2 | 14.8 | 1,901.3 | 7,152.2 |
| TIR/MuDR | | 3.8 | 9.9 | 4,291.2 | 12,449.1 |
| MITE | 0.56 | 465.1 | 1188.4 | 8,138.3 | 8,119.6 |
| Maverick/Maverick | 0.05 | 44.2 | 761.9 | 22,994.6 | 9,898.3 |
| Helitron/Helitron | 0.18 | 2.1 | 10.3 | 726.5 | 9,262.4 |
| LTR/Retrovirus | - | 0.7 | 9.9 | | |
| LINE/R2 | - | 0 | 0.6 | | |
| SINE/7SL (Alu) | - | 1.2 | 0.3 | | |
| TIR/ISL2EU | - | 17.7 | 16.1 | | |
| TIR/Ginger1 | - | 10.2 | 6 | | |
| TIR/Academ | - | 5.1 | 8.5 | | |
| TIR/MuDR | - | 3.8 | 9.9 | | |
| TIR/CACTA | | 1.9 | 0.8 | | |
| TIR/P | - | 0.6 | 0.6 | | |

### S5 The number of retained small RNA molecules after each processing step

| Tissue | RIN value of RNA | Clean reads (after FastX) | After adapter cut (18-40 nt) | riboRNA fragments removed | sRNA molecules (22 nt) | miRNAs (22nt, hit in miRBase) | miRNAs mapped to genes (21-24 nt) | piRNAs (25-30 nt) | Unique piRNAs (25-30 nt) | piRNAs mapped to TEs (25-30 nt) | ping pong occurrence (10 bp overlap) |
|---|---|---|---|---|---|---|---|---|---|---|---|
| Testis#1 | 9.6 | 18,628,420 | 18,387,280 | 1,393,269 | 416,693 | 169,873 | 297,076 | 14,491,636 | 5,844,245 | 3,045,727 | 114,173 |
| Testis#2 | 8.9 | 20,632,085 | 20,127,121 | 702,889 | 743,042 | 306,713 | 360,612 | 14,928,178 | 5,812,644 | 2,443,896 | 90,569 |
| Testis#3 | 9.4 | 15,341,619 | 15,148,223 | 1,731,806 | 343,516 | 128,145 | 244,710 | 11,028,768 | 4,724,341 | 2,319,293 | 86,486 |
| Testis#4 | 9.3 | 18,141,749 | 18,022,557 | 682,282 | 220,515 | 84,628 | 124,381 | 15,672,237 | 2,598,248 | 1,264,088 | 37,855 |
| Ovary#1 | 9.3 | 17,133,153 | 16,652,089 | 2,708,832 | 1,619,086 | 959,868 | 852,065 | 7,423,955 | 2,535,654 | 2,503,814 | 31,837 |
| Ovary#2 | 9.6 | 15,112,304 | 14,382,824 | 2,298,340 | 2,025,800 | 1,191,104 | 601,043 | 5,576,455 | 1,023,454 | 1,805,324 | 3,567 |
| Ovary#3 | 8.7 | 13,323,488 | 11,664,671 | 4,260,265 | 1,226,450 | 504,288 | 482,384 | 2,820,885 | 882,817 | 897,586 | 5,151 |

**S6 The ratios of the summed expression of (A) piRNA pathway genes, (B) NuRD and associated repressive complex genes, and (C) TRIM28 to the summed expression of miRNA pathway genes in species with diverse genome sizes.** Only genes that occurred in all species were included. The species and their genome sizes (in Gb) from left to right are: *Platyplectrum ornatum* (1), *Gallus gallus* (1.3), *Danio rerio* (1.4), *Xenopus tropicalis* (1.7), *Anolis carolinensis* (2.2), *Mus musculus* (2.5), *Geotrypetes seraphini* (3.8), *Rhinatrema bivittatum* (5.3), *Caecilia tentaculate* (5.5), *Pleurodeles waltl* (20), *Ranodon sibiricus* (21), *Ambystoma mexicanum* (32), *Protopterus annectens* (43), *Cynops orientalis* (44), and *Protopterus aethiiopicus* (~130).

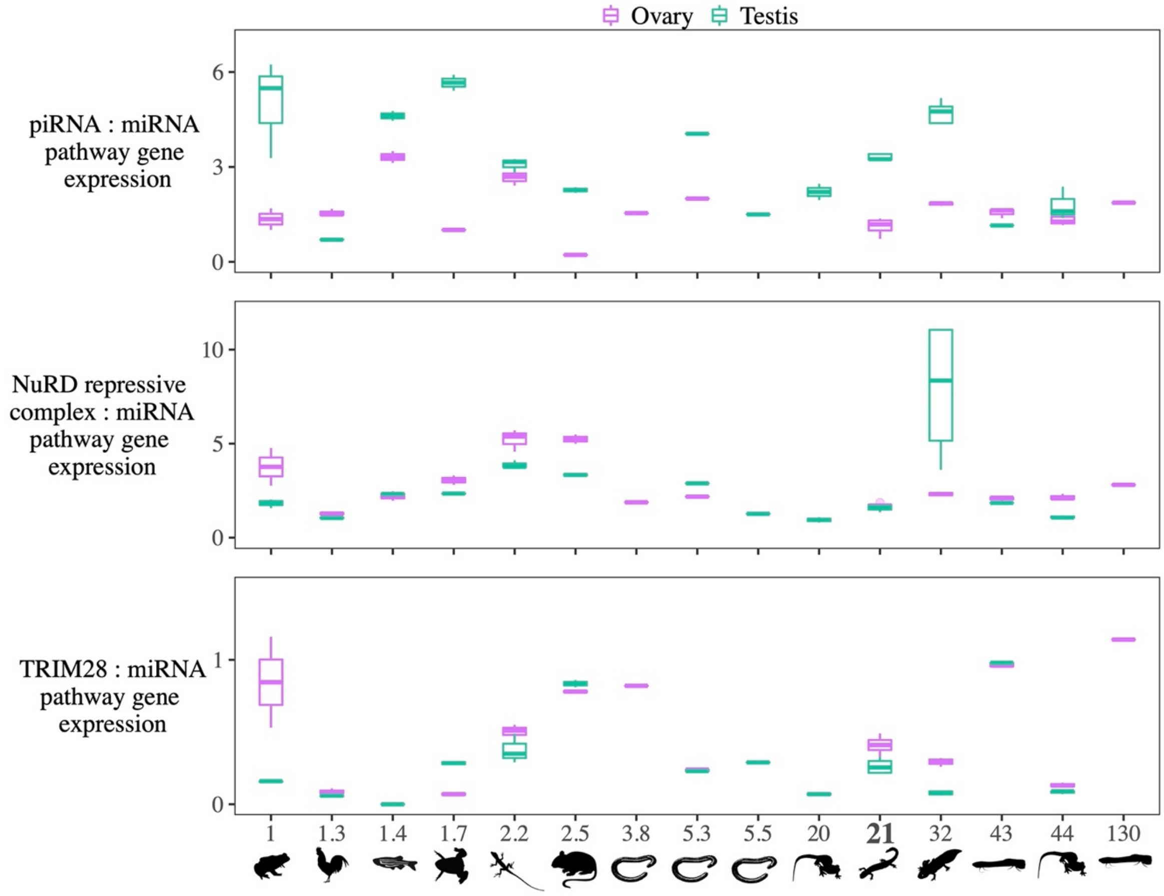

**S7 The summed expression of (A) piRNA pathway genes, (B) NuRD and associated repressive complex genes, (C) TRIM28, and (D) miRNA pathway genes in species with diverse genome sizes measured as TPM.** The species and their genome sizes from left to right are: *Platyplectrum ornatum* (1), *Gallus gallus* (1.3), *Danio rerio* (1.4), *Xenopus tropicalis* (1.7), *Anolis carolinensis* (2.2), *Mus musculus* (2.5), *Geotrypetes seraphini* (3.8), *Rhinatrema bivittatum* (5.3), *Caecilia tentaculate* (5.5), *Pleurodeles waltl* (20), *Ranodon sibiricus* (21), *Ambystoma mexicanum* (32), *Protopterus annectens* (43), *Cynops orientalis* (44), and *Protopterus aethiiopicus* (~130).

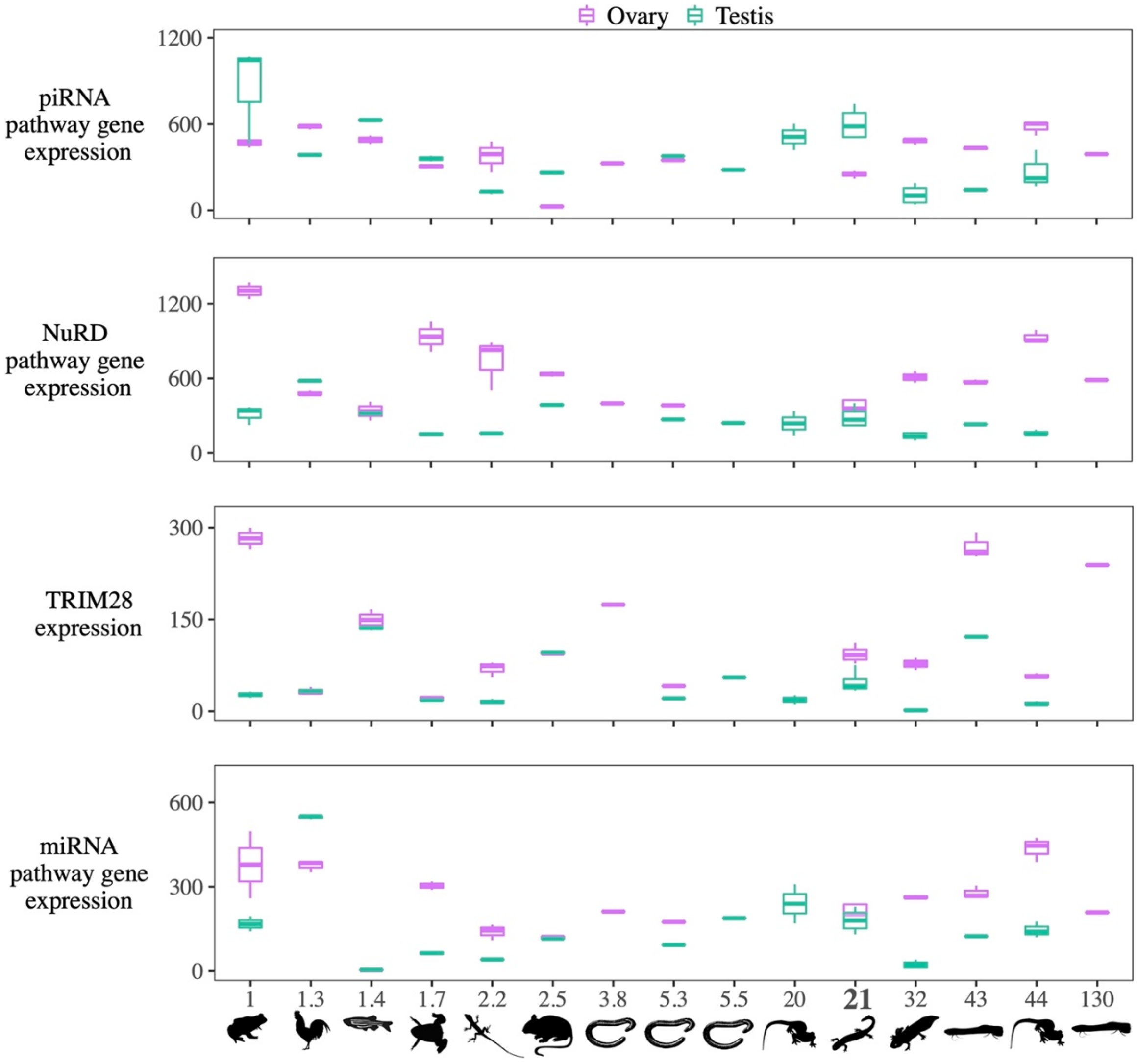

**S8 Samples used for the analysis of piRNA pathway, miRNA pathway, NuRD and associated repressive complex genes, and TRIM28 genes**

| Animal type | Species name | Genome Size: Gb | No. of Testis samples | No. of ovary samples | Number of De novo assembled contigs | N50 of contigs |
|---|---|---|---|---|---|---|
| lungfish | *Protopterus aethiopicus* | 130 | | 1 | 80,874 | 2,584 |
| salamander | *Cynops orientalis* | 44 | 3 | 3 | 287,540 | 1,909 |
| lungfish | *Protopterus annectens* | 43 | 2 | 3 | 191,206 | 2,290 |
| salamander | *Ambystoma mexicanum* | 32 | 4 | 3 | 250,353 | 1,995 |
| salamander | *Ranodon sibiricus* | 21 | 4 | 4 | 510,439 | 1,250 |
| salamander | *Pleurodeles waltl* | 20 | 2 | | 303,910 | 2,232 |
| caecilian | *Caecilia tentaculata* | 5.5 | 1 | | 129,038 | 1,790 |
| caecilian | *Rhinatrema bivittatum* | 5.3 | 1 | 1 | 542,424 | 1,119 |
| caecilian | *Geotrypetes seraphini* | 3.8 | | 1 | 334,360 | 1,527 |
| mouse | *Mus musculus* | 2.5 | 2 | 2 | 327,400 | 983 |
| lizard | *Anolis carolinensis* | 2.2 | 3 | 3 | 262,759 | 1,040 |
| frog | *Xenopus tropicalis* | 1.7 | 2 | 2 | 246,914 | 1,068 |
| fish | *Danio rerio* | 1.4 | 2 | 2 | 168,654 | 1,757 |
| bird | *Gallus gallus* | 1.3 | 3 | 3 | 757,972 | 2,154 |
| frog | *Platyplectrum ornatum* | 1.07 | 3 | 2 | 367,023 | 1,370 |

**S8 Samples used for the analysis of piRNA pathway, miRNA pathway, NuRD and associated repressive complex genes, and TRIM28 genes**

| Species | Individuals | Code of dataset | Sequencing strategy | Total length of clean paired reads |
|---|---|---|---|---|
| *Platyplectrum ornatum* | Female_F1 | SRR13734432 | PE126 | 6,884,372,718 |
| *Platyplectrum ornatum* | male_M1 | SRR13734436 | PE126 | 3,519,736,155 |
| *Platyplectrum ornatum* | female_F2 | SRR13734440 | PE126 | 3,684,200,728 |
| *Platyplectrum ornatum* | male_M2 | SRR13734445 | PE126 | 3,159,099,352 |
| *Platyplectrum ornatum* | male_M3 | SRR13734453 | PE126 | 5,591,344,200 |
| *Gallus gallus* | M1 | SRR21413816 | PE150 | 6,801,083,731 |
| *Gallus gallus* | M2 | SRR21413817 | PE150 | 6,570,577,470 |
| *Gallus gallus* | M3 | SRR21413818 | PE150 | 7,021,840,670 |
| *Gallus gallus* | FM1 | SRR21413813 | PE150 | 6,709,497,312 |
| *Gallus gallus* | FM2 | SRR21413814 | PE150 | 6,570,023,137 |
| *Gallus gallus* | FM3 | SRR21413815 | PE150 | 6,800,803,540 |
| *Danio rerio* | M1 | SRR6841469<br>SRR6841470<br>SRR6841471 | PE76 | 14,132,566,824 |
| *Danio rerio* | M2 | SRR6841472<br>SRR6841473<br>SRR6841474 | PE76 | 12,451,819,259 |
| *Danio rerio* | FM1 | SRR6841457<br>SRR6841458<br>SRR6841459 | PE76 | 15,041,093,899 |
| *Danio rerio* | FM2 | SRR6841460<br>SRR6841461<br>SRR6841462 | PE76 | 14,383,695,630 |
| *Xenopus tropicalis* | M1 | SRR5412279 | SR101 | 4,491,825,839 |
| *Xenopus tropicalis* | M2 | SRR5412280 | SR101 | 4,959,481,791 |
| *Xenopus tropicalis* | FM1 | SRR5412277 | SR101 | 3,522,394,964 |
| *Xenopus tropicalis* | FM2 | SRR5412278 | SR101 | 3,433,990,644 |
| *Anolis carolinensis* | M1 | SRR5412171 | SR101 | 2,037,335,955 |
| *Anolis carolinensis* | M2 | SRR5412172 | SR101 | 2,328,660,042 |
| *Anolis carolinensis* | M3 | SRR5412173 | SE90 | 3,429,516,547 |
| *Anolis carolinensis* | FM1 | SRR5412168 | SR101 | 2,473,892,259 |
| *Anolis carolinensis* | FM2 | SRR5412169 | SR101 | 2,352,520,907 |
| *Anolis carolinensis* | FM3 | SRR5412170 | SE90 | 3,531,532,320 |
| *Mus musculus* | M1 | SRR5412203 | SR101 | 1,955,779,276 |

| | | | | |
|---|---|---|---|---|
| *Mus musculus* | M2 | SRR5412204 | SR101 | 3,318,896,675 |
| *Mus musculus* | FM1 | SRR5412201 | SR101 | 2,277,704,981 |
| *Mus musculus* | FM2 | SRR5412202 | SR101 | 2,023,556,350 |
| *Rhinatrema bivittatum* | M1 | SRR5591432 | PE101 | 5,014,521,076 |
| *Rhinatrema bivittatum* | FM1 | ERR3132330 | PE150 | 22,243,030,188 |
| *Ambystoma mexicanum* | M1 | SRR2885287 | PE100 | 3,557,193,043 |
| *Ambystoma mexicanum* | M2 | SRR2885288 | PE100 | 3,018,140,755 |
| *Ambystoma mexicanum* | M3 | SRR2885289 | PE100 | 6,947,066,098 |
| *Ambystoma mexicanum* | M4 | SRR2885290 | PE100 | 2,254,388,959 |
| *Ambystoma mexicanum* | FM1 | SRR2885284 | PE100 | 3,046,610,392 |
| *Ambystoma mexicanum* | FM2 | SRR2885285 | PE100 | 2,480,135,684 |
| *Ambystoma mexicanum* | FM3 | SRR2885286 | PE100 | 2,806,064,782 |
| *Protopterus annectens* | M1 | SRR2028017 | PE90 | 7,004,563,990 |
| *Protopterus annectens* | FM1 | SRR2027978 | PE90 | 6,124,764,166 |
| *Protopterus annectens* | FM2 | SRR2027979 | PE90 | 6,518,099,880 |
| *Protopterus annectens* | FM3 | SRR2027980 | PE90 | 6,494,872,072 |
| *Protopterus aethiopicus* | FM1 | SRR7240708 | PE150 | 6,493,306,793 |
| *Cynops orientalis* | Ovary3_FG3 | SRR10305546 | PE122 | 4,994,921,293 |
| *Cynops orientalis* | Ovary2_FG2 | SRR10305545 | PE122 | 5,326,272,623 |
| *Cynops orientalis* | Ovary1_FG1 | SRR10305544 | PE122 | 4,550,235,130 |
| *Cynops orientalis* | Testis3_M3 | SRR10305543 | PE122 | 5,720,828,649 |
| *Cynops orientalis* | Testis2_M2 | SRR10305542 | PE122 | 5,634,805,471 |
| *Cynops orientalis* | Testis1_M1 | SRR10305541 | PE122 | 4,972,425,431 |
| *Ranodon sibiricus* | Testis_WQ05 | CRR610083 | PE150 | 4,468,359,749 |
| *Ranodon sibiricus* | Testis_WQ07 | CRR610084 | PE150 | 5,354,698,049 |
| *Ranodon sibiricus* | Testis_WQ08 | CRR610085 | PE150 | 5,054,882,981 |
| *Ranodon sibiricus* | Testis_WQ10 | CRR610086 | PE150 | 4,940,685,042 |
| *Ranodon sibiricus* | Ovary_WQ02 | CRR610079 | PE150 | 5,771,046,105 |
| *Ranodon sibiricus* | Ovary_WQ03 | CRR610080 | PE150 | 5,466,338,283 |
| *Ranodon sibiricus* | Ovary_WQ04 | CRR610081 | PE150 | 5,716,393,060 |
| *Ranodon sibiricus* | Ovary_WQ06 | CRR610082 | PE150 | 5,612,656,781 |
| *Pleurodeles waltl* | M3 | DRR138635 | PE106 | 3,580,009,573 |
| *Pleurodeles waltl* | M4 | DRR138637 | PE106 | 3,397,990,304 |
| *Caecilia tentaculata* | Testis | SRR5591445 | PE101 | 5,326,917,714 |
| *Geotrypetes seraphini* | Ovary | ERR3849999 | PE150 | 27,001,609,155 |
